# A distant global control region is essential for normal expression of anterior *HOXA* genes during mouse and human craniofacial development

**DOI:** 10.1101/2022.03.10.483852

**Authors:** Andrea Wilderman, Eva D’haene, Machteld Baetens, Tara N. Yankee, Emma Wentworth Winchester, Nicole Glidden, Ellen Roets, Jo Van Dorpe, Sarah Vergult, Timothy C. Cox, Justin Cotney

## Abstract

Defects in embryonic patterning resulting in craniofacial abnormalities account for approximately 1/3 of birth defects. The regulatory programs that build and shape the face require precisely controlled spatiotemporal gene expression, achieved through tissue-specific enhancers. Large regions with coactivation of enhancer elements and co-regulation of multiple genes, referred to as superenhancers, are important in determining cell identity and perturbation could result in developmental defects. Building upon a previously published epigenomic atlas of human embryonic craniofacial tissue in which we identified over 75,000 putative embryonic craniofacial enhancer regions, we have identified 531 superenhancer regions unique to embryonic craniofacial tissue, including 37 which fall in completely noncoding regions. To demonstrate the utility of this data for the understanding of craniofacial development and the etiology of craniofacial abnormalities, we focused on a craniofacial-specific superenhancer in a ∼600kb noncoding region located between *NPVF* and *NFE2L3*. This region harbors over 100 individual putative craniofacial enhancer segments and 7 *in vivo* validated craniofacial enhancers from primary craniofacial tissue as well as strong enhancer activation signatures in a culture model of cranial neural crest cell (CNCC) development. However, none of the directly adjacent genes have been implicated in neural crest specification, craniofacial development, or abnormalities. To identify potential regulatory targets of this superenhancer region, we characterized three-dimensional chromatin structure of this region in CNCCs and mouse embryonic craniofacial tissues using multiple techniques (4C-Seq, HiC). We identified long range interactions that exclude most intervening genes and specifically target the anterior portion of the *HOXA* gene cluster located 1.2 to 1.8 Mb away. We demonstrate the specificity of the enhancer region for regulation of anterior *HOXA* genes through CRISPR/Cas9 editing of human embryonic stem cells. Mice homozygous for deletion of the superenhancer confirm the specificity of the enhancer region and demonstrate that the region is essential for viability. At fetal stages homozygotes develop at the same rate as heterozygous and wild type littermates but die at P0-P1 and have high penetrance of orofacial clefts that phenocopy previously described *Hoxa2-/-* mice. Moreover, we identified a *de novo* deletion partially overlapping the superenhancer in a human fetus with severe craniofacial abnormalities. This evidence suggests we have identified a critical noncoding locus control region that specifically regulates anterior *HOXA* genes and whose deletion is likely pathogenic in human patients.

## Introduction

Proper control of gene expression during development and in adult tissues is achieved in part through regulatory sequences typically referred to as enhancers. Enhancers are collections of transcription factor binding sites that have been shown to control gene expression in a temporal and tissue-specific manner (Spitz and Furlong, 2012; Fulco et al., 2016). Some genes are regulated by a single enhancer in a particular tissue or context (Lettice et al., 2003), but most genes are regulated by multiple enhancers with each contributing to a portion of the overall target gene expression (Osterwalder et al., 2018). Coactivated clusters of individual enhancer elements and co-regulation of multiple nearby genes has been seen as a strong biomarker for cell tissue identity genes. These activated regulatory landscapes, referred to as superenhancers, are frequently associated with genes for cell-type specific transcription factors, giving them an important role in determining cell identity (Whyte et al., 2013; Hnisz et al., 2013). Due to their size, sequence composition, and largely unknown contributions to specific gene regulation, the impact of noncoding mutations and copy number variation in superenhancers presents a complicated area of study. Thus far copy number changes within superenhancer regions have been associated with tumorigenesis (Hnisz et al., 2013), and disease-associated SNPs are enriched in superenhancers active in cell types relevant to the disease (Huang et al., 2018). While superenhancer regions are also potential regulators of early developmental processes and their role in developmental defects has yet to be clearly established. Isolated clinical reports have indicated that insertion of a superenhancer into a new context can result in craniofacial abnormalities (Middelkamp et al., 2019) but how frequently this occurs is unknown.

Superenhancers are identified via active chromatin marks combined with a high degree of occupancy by master regulators including the Mediator complex (Whyte et al., 2013; Khan and Zhang, 2016). Whether superenhancers constitute a unique paradigm for gene regulation has been a question since their definition (Pott and Lieb, 2015; Blobel et al., 2021; Hay et al., 2016). The behavior of enhancer elements within superenhancers are varied. Studies suggest that individual enhancer elements within superenhancers cooperate in an additive, redundant or synergistic manner (Hay et al., 2016; Osterwalder et al., 2018; Moorthy et al., 2017; Guerrero et al., 2010; Choi et al., 2021). Other studies have characterized superenhancers in which a single potent enhancer element drives the majority of the effect (Moorthy et al., 2017; Farooq et al., 2021). Interestingly the potency of a superenhancer or the individual enhancer elements within it cannot be definitively predicted by degree of conservation (Ahituv et al., 2007; Nolte et al., 2014; Osterwalder et al., 2018; Dickel et al., 2018).

The major challenges in the study of superenhancers are twofold. First, their specificity requires that superenhancers be identified in the relevant tissue at the developmental stage of interest. Second, perturbation of a superenhancer and downstream consequences may only be apparent during developmental or specific conditions that are difficult to create experimentally (Dave et al., 2017; Saravanan et al., 2020). Despite functional annotations of active chromatin states being available for numerous human tissues, early developmental stages are underrepresented and therefore many superenhancers y are unknown.

We previously used ChIP-seq for six histone modifications coupled with imputation of additional epigenomic characteristics to provide epigenomic annotations during critical stages of human craniofacial development (Wilderman et al., 2018). This approach revealed that variants associated with common nonsyndromic craniofacial abnormalities such as nonsyndromic cleft lip and palate (NSCL/P) are enriched in enhancers active in early developmental stages. We and others have subsequently shown with this data that variants associated with normal facial variation are enriched in enhancers active in later developmental stages (Wilderman et al., 2018; White et al., 2021; Bonfante et al., 2021). The comprehensive nature of the data obtained from our previous investigation makes it ideally suited to the identification of craniofacial-specific superenhancers and investigation of their role in nonsyndromic craniofacial malformations.

Here we report novel human craniofacial-specific superenhancer regions in developing craniofacial tissue spanning organogenesis and describe their general characteristics. Additionally, we identified craniofacial-specific superenhancers that did not harbor a known gene and tested the function of one such region. Our examination identified a novel superenhancer region that interacts with the *HOXA* locus in human and mouse embryonic craniofacial tissues. We demonstrate that deletion of this novel superenhancer in mice decreases anterior *Hoxa* gene expression in pharyngeal arch tissue and recapitulates the distinct craniofacial phenotypes reported in *Hoxa2* null mice. We include discussion of patients with deletions overlapping this region and the potential for pathogenicity of noncoding mutations in this region in humans.

## Results

### Identification of novel craniofacial superenhancers from epigenomic atlas

We reasoned that superenhancers active during craniofacial development might be enriched for novel master regulator genes as well as regions of the genome likely to be linked to craniofacial abnormalities. To address this question we first sought to identify superenhancer regions in a systematic fashion across craniofacial development. Using 75,928 previously identified craniofacial enhancer segments and H3K27ac ChIP-seq signals, (Wilderman et al., 2018) we identified superenhancer regions using the ROSE algorithm (Methods, (Whyte et al., 2013), **Figure 1a**). From 17 samples of human craniofacial tissue across five embryonic and one fetal stage encompassing the major events in craniofacial development, we identified an average of 1861 superenhancers per sample. These represent 4,339 distinct superenhancer regions across the developmental trajectory (**Supplemental Table 1**).

**Figure 1.**
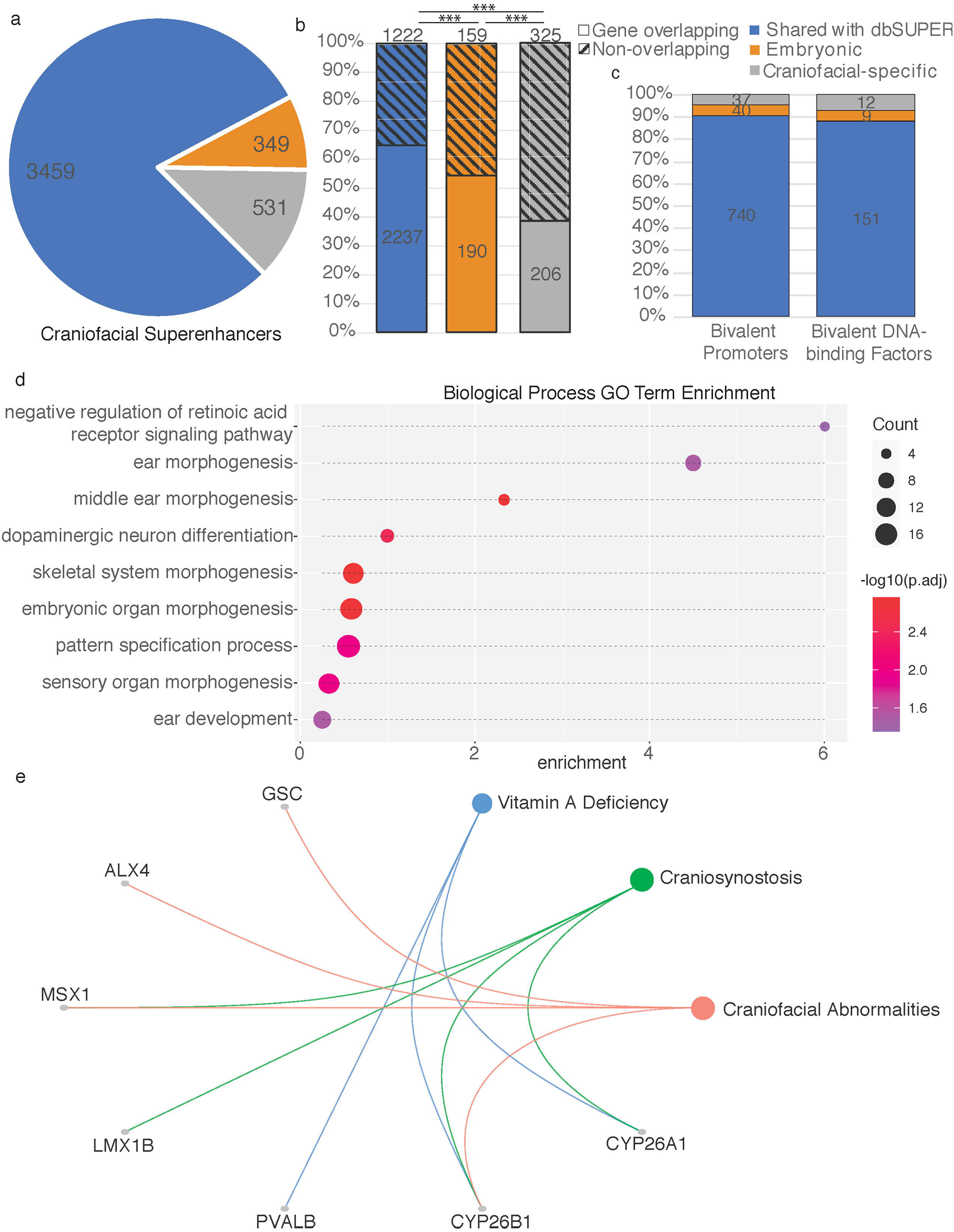
Characteristics of human embryonic craniofacial superenhancers relate to specialized developmental functions. a. Sharing of superenhancer regions (see Methods) with tissues and cell types within dbSUPER (blue), or only with human embryonic heart (termed Embryonic enhancers, orange). Those unique to the human embryonic craniofacial tissue are shown in gray. b. Percentages of shared or unique superenhancer regions which encompass a TSS (Gene-overlapping) or not. The Fisher Exact Test was used to compare the proportions of gene overlapping and non-overlapping superenhancers, *** p<0.001. c. Summary of TSS encompassed in each category of superenhancer regions that correlate with genes previously identified to have bivalent promoters, and a further subset of transcription factors with bivalent promoters. d. Gene Ontology terms enriched in genes for which the TSS is encompassed by superenhancer regions unique to craniofacial tissue. e. Disease Ontology relationships with genes for which the TSS encompassed by superenhancer regions unique to craniofacial tissue are among the previously determined genes with bivalent promoters.

Superenhancers typically encompass their tissue-specific targets or reside in the introns of their target genes (Whyte et al., 2013; Hnisz et al., 2013). Consistent with this, approximately 90% of individual craniofacial superenhancers overlap at least one gene (average 2.3+/-0.22 protein-coding genes per embryonic sample, Mean +/-SD). We found similar results for superehancers identified in tissues profiled by the Roadmap Epigenomics and ENCODE projects, which were retrieved from dbSuper (Khan and Zhang, 2016) (**Figure 1 Supplement 1**; **Supplemental Table 2**). These embryonic craniofacial superenhancers often overlapped bivalent promoters of DNA binding factors, including the homeobox transcription factors (Wilderman et al., 2018). Genes encompassed by these craniofacial superenhancers were also systematically enriched for craniofacial disease related ontologies including craniofacial abnormalities, abnormalities of the midface, abnormalities of eyes (e.g. exophthalmos), and terms for facial characteristics (e.g. frontal bossing, pointed chin, round face) (**Supplemental Table 3**)

Having demonstrated that our craniofacial superenhancer calls were consistent in both size and scale to other human tissues and were enriched for craniofacial relevant biology, we next investigated whether any superenhancers were truly specific to craniofacial development. To achieve this we determined intersections of the 4,339 distinct craniofacial superenhancer regions with all superenhancer calls from dbSuper as well as the human embryonic heart (VanOudenhove et al., 2020) **(Figure 1**). In total we identified 531 superehancer regions that were uniquely superenhancer calls in craniofacial tissue. These regions tended to be slightly smaller than the distribution of all craniofacial superenhancers (median 37.4kb vs 52.6kb) and tended to contain fewer genes (**Supplemental Table 4**). More than half of the craniofacial specific superenhancers did not encompass the transcription start site (TSS) of a gene (see **Figure 1b**). Craniofacial specific superenhancers that did encompass one or more TSS did so for genes enriched for functions related to development, including embryonic organ development, skeletal system morphogenesis and sensory organ morphogenesis including morphogenesis of the middle ear, reflecting the larger craniofacial superenhancer set and suggesting additional specialized activity (**Figure 1d**). Furthermore, a subset of these genes were previously determined to have bivalent promoters in human embryonic craniofacial tissue (**Figure 1c**) and are implicated in craniofacial abnormalities (**Figure 1e**).

When we examined those superenhancers that did not overlap a TSS, the genes flanking these superenhancers (n=584 genes, n=325 regions) were enriched for Biological Process Gene Ontology terms related to embryonic kidney development, (**Figure 1 Supplement 2**). However, the genes were enriched for Disease Ontology terms related to craniofacial abnormalities, clefting as well as types of tumors considered neurocristopathies (Etchevers et al., 2006) **(Figure 1 Supplement 3**).

We thus reasoned that craniofacial specific superenhancers that are located in large noncoding regions or “gene deserts” might be novel regions important for craniofacial development. To prioritize these lists of craniofacial specific superenhancers for regions that could be experimentally studied we first examined individual enhancers that had been tested *in vivo* and demonstrated craniofacial activity in the developing mouse.

### Gene Desert on Chromosome 7 Contains Superenhancer Regions Unique to Embryonic Craniofacial Tissue

Of the craniofacial-specific superenhancers located entirely in noncoding regions, 152 contained at least 1 segment that had been tested for activity in the VISTA enhancer database. Of these, 37 contained at least one enhancer with validated craniofacial activity (**Supplemental Table 5**). The region containing the greatest number of confirmed craniofacial enhancers was a 600kb span of chromosome 7. This gene desert (chr7:25,400,000-26,000,000; hg19) resides between *NPVF* and *MIR148A* (**Figure 2a**) and contains embryonic craniofacial-specific superenhancer regions with validated craniofacial enhancer function. The superenhancer regions between chr7:25,580,400-25,849,400 are unique to human embryonic craniofacial tissue, not having been identified as such in human embryonic heart tissue (VanOudenhove et al., 2020) or any of the 102 human tissues and cell lines analyzed by dbSuper (Khan and Zhang, 2016) and https://asntech.org/dbsuper/index.php) **(Figure 2b)**. We identified six enhancer segments tested by the VISTA Enhancer Browser (https://enhancer.lbl.gov/; (Visel et al., 2007) that drove reporter expression in mouse craniofacial tissue at E11.5. An additional human enhancer segment, HACNS50 (Prabhakar et al., 2008) was also tested and found to drive strong reporter expression in mouse embryonic craniofacial and limb tissue. **(Figure 2c)**.

**Figure 2.**
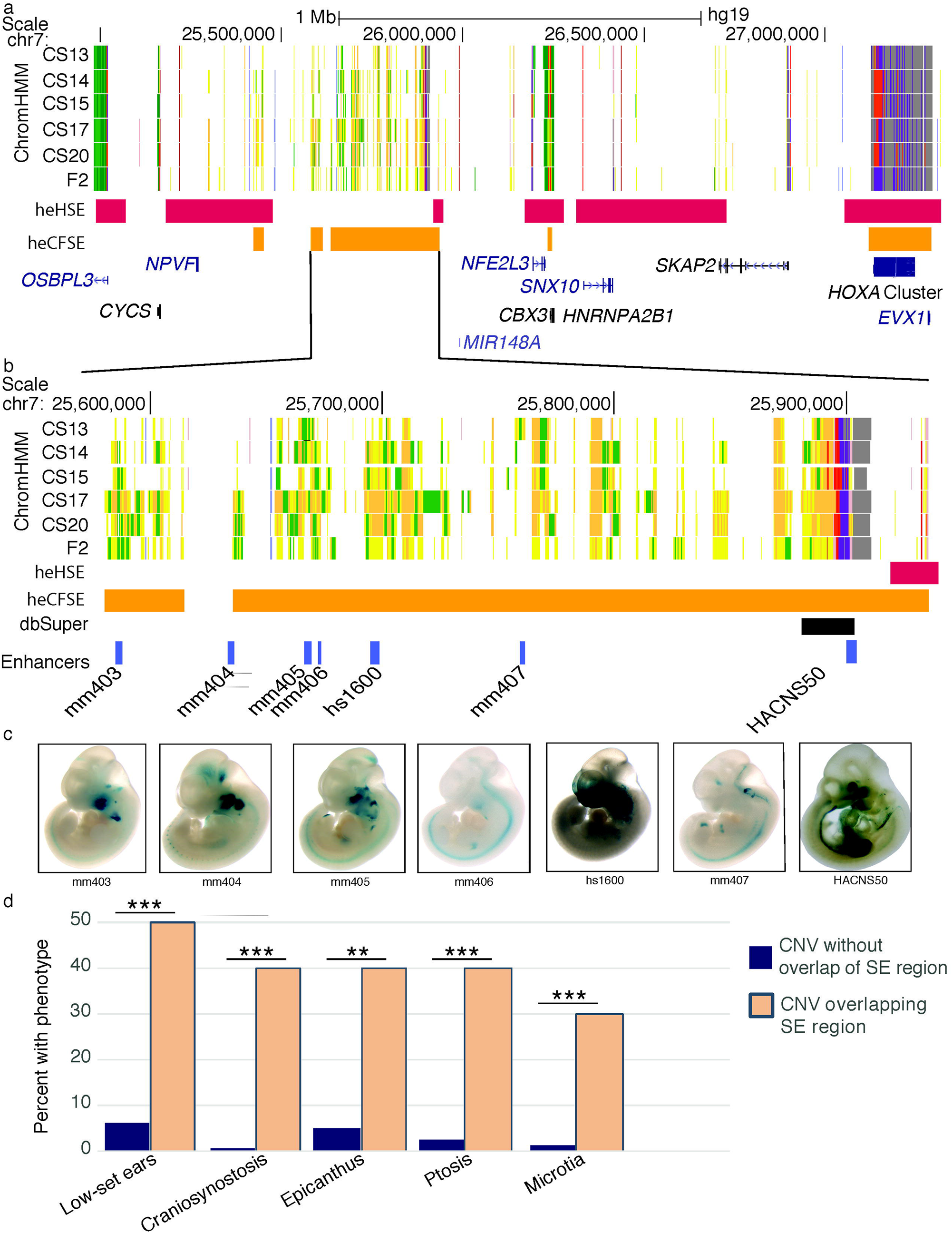
Location and functional characterization of a putative novel craniofacial superenhancer. a. Region of human chromosome 7 containing a large 500kb window lacking any annotated protein-coding genes with extensive enrichment of activated enhancer (yellow and orange) and transcriptionally active (green) segment annotations in human craniofacial tissue. CS (Carnegie stage). Locations of human embryonic craniofacial superenhancers (heCFSE) are represented by orange bars, human embryonic heart superenhancers (heHSE) by dark pink bars and superenhancers found in the dbSuper database by black bars. b. Enlargement of two superenhancers with multiple validated craniofacial enhancer segments. Enhancers with mm or hs designations were identified through the Vista Enhancer Browser (c). In this study we tested and validated the craniofacial enhancer activity of HACNS50, located within the bivalent chromatin state at the right of the enlargement. d. Frequency of phenotypes present in ten individuals within the DECIPHER Database with CNVs overlapping chr7:25,580,400-25,849,400 (hg19) and phenotype descriptions compared to frequency of those phenotypes in the DECIPHER Database for CNVs not overlapping the region. Statistical test is the Fisher Exact Test, ** p<0.01, *** p<0.001.

We then turned to a curated database of patients with a variety of developmental abnormalities likely due to copy number variation. Of the 21 Individuals within the DECIPHER Database (deciphergenomics.org; Firth et al., 2009) with copy number variants (CNVs) overlapping the region (chr7:25,580,400-25,849,400;hg19), ten had a single CNV which could be considered causastive. Those 10 individuals demonstrated a higher incidence of craniofacial abnormalities than the database as a whole (**Figure 2d; Figure 2 Supplement 1; Figure 2 Supplement 2; Supplemental Table 6**). Together these findings strongly suggest that this superenhancer region is specific for craniofacial development and associated with craniofacial abnormalities when copy number is altered. We therefore focused on this region to identify the critical genes it might regulate and what role it might play in craniofacial development.

To identify target genes which might be regulated by this craniofacial-specific region, we looked for craniofacially-relevant genes located nearby. When considering genes up to 500kb in either direction of the gene desert, we observed *NPVF, MIR28A, OSBPL3, CYCS, C7ORF31, NFE2L3, HNRNPA2B1, CBX3* and *SNX10*, none of which have been specifically associated with craniofacial development or defects (Lee et al., 2017; Braconi et al., 2010; (Li et al., 2016). When we examined expression of all these genes in primary human craniofacial tissues and all GTEX tissues, we found similar levels across most tissues (**Figure 2 Supplement 3**; (Yankee et al., 2022). Only *SNX10* had elevated specificity of expression but was most highly expressed in the adult brain. These findings raised the possibility that either one of these genes plays an unappreciated role in craniofacial development or the target of this superenhancer may lie a considerable distance away.

The region being considered was then expanded. This revealed *SKAP2* and the *HOXA* gene cluster approximately 1.5Mb downstream. *SKAP2* has not previously been implicated in craniofacial development. In contrast, *HOXA* genes have been linked to a number of syndromes that include craniofacial abnormalities in both mouse and human (Alasti et al., 2008; Barrow and Capecchi, 1999; Quinonez and Innis, 2014; Tischfield et al., 2005)). Despite the distance of the *HOXA* cluster from the superenhancer, its tissue relevance indicated these genes as feasible targets.

### Chromatin architecture of chr7 25,000,000-28,000,000 suggests the superenhancer makes long-distance contacts with the *HOXA* cluster

To determine if this craniofacial specific superenhancer region could indeed target the *HOXA* gene cluster located >1MB downstream, we first examined publicly available HiC data from a variety of cell types. In human embryonic stem cells (hESCs) we found that the entire superenhancer region formed a topologically associated domain (TAD). However, it was unclear if this region formed longer range interactions that incorporated the *HOXA* cluster (**Figure 3a**). Other adult tissues and cell types showed similar trends (**Figure 3 Supplement 1**), but this region was specifically identified as active in human craniofacial development, leaving open the possibility that interactions might only happen in craniofacial relevant contexts. We therefore performed chromosome conformation capture experiments (HiC) in both primary human embryonic craniofacial tissue and a recently described culture model of differentiated cranial neural crest cells (CNCC) (Leung et al., 2016).

**Figure 3.**
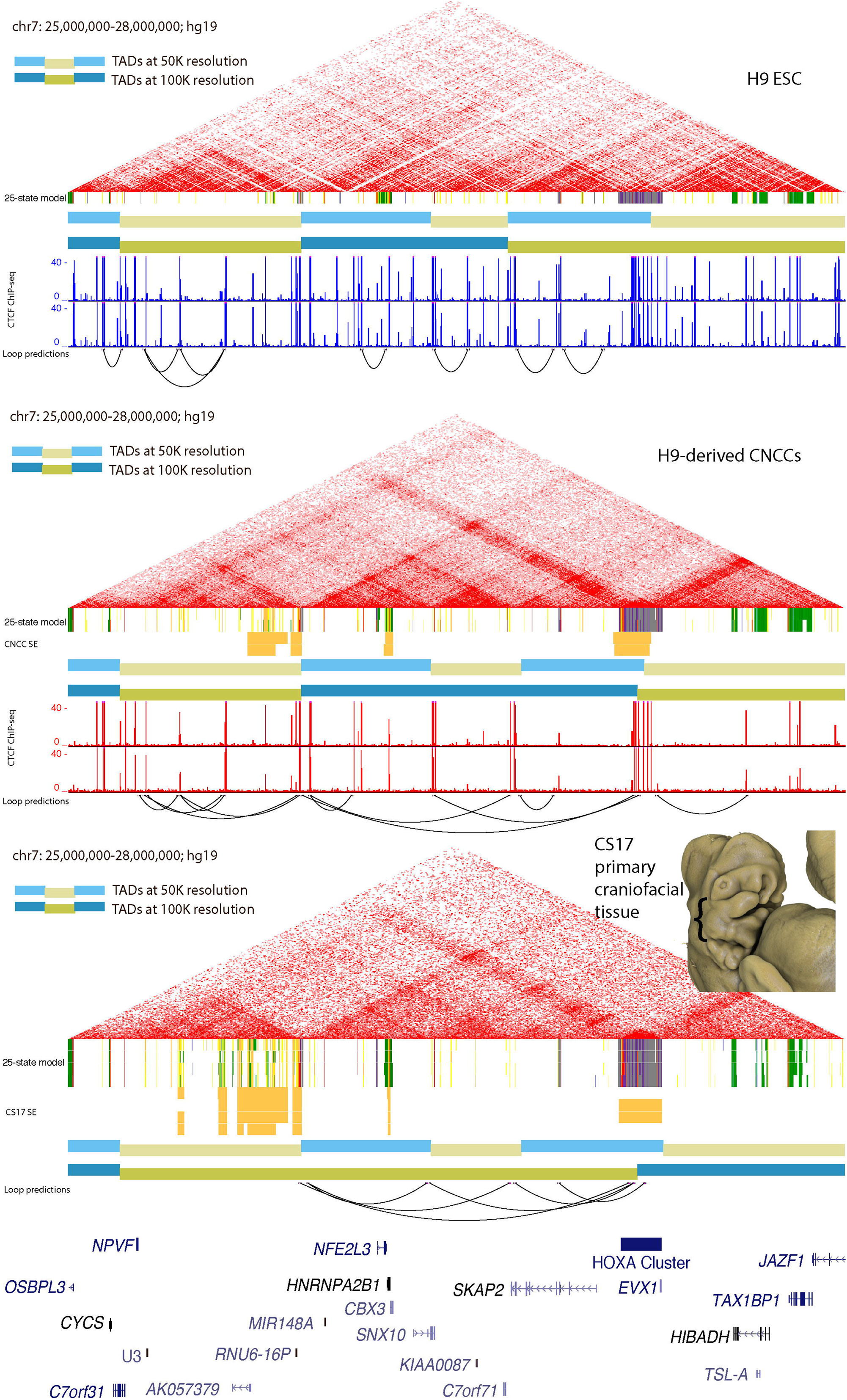
Chromatin Architecture in Primary Human Embryonic Craniofacial Tissue Suggests Interaction between HOXA Gene Cluster and Gene Desert Superenhancer on Chromosome 7. HiC of H9 human embryonic stem cells (hESCs)(a), cranial neural crest cells (CNCCs) derived from H9 hESCs (b) and CS17 primary human embryonic craniofacial tissue(c). TADs called at two different resolutions, 50kb (light blue/light yellow) and 100kb (blue/dark yellow). Superenhancers (for CNCCs and CS17 tissue) determined by ROSE algorithm. CTCF ChIP-seq data from (Long et al. 2020; GSE145327). ChromHMM chromatin states from 25-state model for H9 and H9-derived CNCCs are shown below their respective HiC interaction plots and chromatin states for CS13-CS20 and F2 human craniofacial tissue are shown below the HiC interaction plot for CS17 primary craniofacial tissue. Individual enhancer segments are yellow and orange. Inset image: 3D rendered Carnegie stage 17 human embryo demonstrate representative staging of tissue used in HiC experiments The embryo was imaged using High Resolution Episcopic Microscopy (HREM): raw data courtesy of Dr Tim Mohun (Francis Crick Institute, London, UK) and provided by the Deciphering the Mechanisms of Developmental Disorders (DMDD) program (https://dmdd.org.uk/).

Consistent with this idea, we found in CNCCs that DNA near the 3’ most boundary region of the TAD identified above formed qualitatively stronger interactions with the *HOXA* gene cluster than hESCs (**Figure 3b**). However, this TAD did not coalesce into a larger domain including those genes in our analysis. Surprisingly, when we analyzed HiC data from primary CS17 (Carnegie stage 17) human embryonic craniofacial samples, a much larger TAD was discovered. This TAD stretched from the 5’-most boundary of the gene desert all the way to the midpoint of the *HOXA* gene cluster (**Figure 3c**). When we inspected gene expression in each of these cell types and tissues, the changes in configuration relative to hESCs coincided with increased expression of *HOXA* genes (**Supplemental Table 7)**.

The proximity to the *HOXA* gene cluster in three-dimensional space confirmed the potential for a regulatory relationship between some or all of the enhancers within the superenhancer and promoters of the anterior *HOXA* genes. *HOXA1* or *HOXA2* are prime candidates as they have gene expression patterns that include craniofacial tissues, and they play an essential role in patterning the cartilage and bone of the skull (Hunt et al., 1991; Gendron-Maguire et al., 1993; (Minoux and Rijli, 2010). Moreover, loss-of-function mutations in these two genes can disrupt normal human craniofacial development. For instance *HOXA1 is* implicated in Bosley-Salih-Alorainy syndrome/Athabaskan brainstem dysgenesis syndrome (OMIM #601536) and *HOXA2*-associated autosomal recessive microtia (OMIM #612290) (Quinonez and Innis, 2014; Alasti et al., 2008; Tischfield et al., 2005). While the regulatory architecture of this gene cluster has been explored with respect to limb development (Kmita et al., 2005; Berlivet et al., 2013; Woltering et al., 2014), the full extent of regulatory elements influencing *HOXA* gene expression in other tissues is unknown.

### Inversion of superenhancer sub-TAD identifies TAD boundary as strong organizing center

The three-dimensional chromatin structure and associated gene expression suggested that the putative enhancer region is part of the *HOXA* regulatory landscape (Bolt and Duboule, 2020; Spitz et al., 2003). To determine if this region is indeed important for anterior *HOXA* gene expression we set out to remove this region from the genome and examine its effects on gene expression in hESCs and CNCCs. Removal of small enhancer modules has frequently resulted in minimal effects on gene expression and mice with very mild phenotypes (Ahituv et al., 2007; Attanasio et al., 2013). Even the disruption of large gene deserts has been reported to result in mice that are overtly normal (Nóbrega et al., 2004; (Dave et al., 2017; Moorthy et al., 2017). Disruption of TAD boundaries has been shown to result in ectopic expression of genes in the newly formed TADs (Lupiáñez et al., 2015; Franke et al., 2016; Hnisz et al., 2016; Weischenfeldt et al., 2017), but the role enhancers play in gene expression in such a scenario is difficult to discern. Here we wanted to study the role of the superenhancer region and if possible separate this from higher order chromatin architecture. We hypothesized that removal of the entire TAD containing the superenhancer might not create new TADs or alter larger chromosome architecture and allow us to identify the specific regulatory outputs of this region. To achieve such a scenario we aimed to identify sites for cutting by Cas9 that were outside both boundaries of the TAD. Current models of chromosome organization indicate that CTCF is an important component for loop formation and TAD boundary establishment (Dixon et al., 2012; Rao et al., 2014; Khoury et al., 2020). When we inspected CTCF binding in a similar model of human neural crest differentiation (Long et al., 2020) we found that a relatively small number of predicted CTCF binding sites in this gene desert were occupied. A single strong CTCF binding event was identified at the 5’-most end of the TAD identified in CNCCs above. At the 3’-most end of this same TAD we observed several strong, closely-spaced CTCF binding sites with the same motif orientation. Consistent with the well-documented insulating role of CTCF, these closely spaced CTCF binding sites directly coincided with the boundary between strongly active and strongly repressed chromatin signals in both CNCCs and primary craniofacial tissues (**Figure 4b**). Having identified more precise boundaries of this putative regulatory domain, we designed guide RNAs to target Cas9 to this region. On the 5’ side of the TAD we selected a sequence downstream of *NPVF* but upstream of the 5’-most CTCF bound site near the TAD boundary. On the 3’ side of the TAD we selected a sequence downstream of the 3’-most CTCF bound site at the TAD boundary but upstream of another cluster of strongly bound CTCF not predicted to be part of this TAD. We then transfected plasmids expressing these guides and Cas9 protein into H9 ESCs and, using PCR, screened for clones that had deleted this region (**Figure 4a; Figure 4 Supplement 1**). Several clones were identified that harbored heterozygous loss of this region but we were unable to remove both alleles of this region. Recent attempts to remove only the 3’ TAD boundary region in hematopoietic stem and progenitor cells (HSPCs) (Zhang et al., 2020) were only able to achieve approximately 50% efficiency and were not assessed for homozygous deletion. These results suggest that this region as a whole is critical for survival in human stem cells following electroporation or other harsh manipulation.

**Figure 4.**
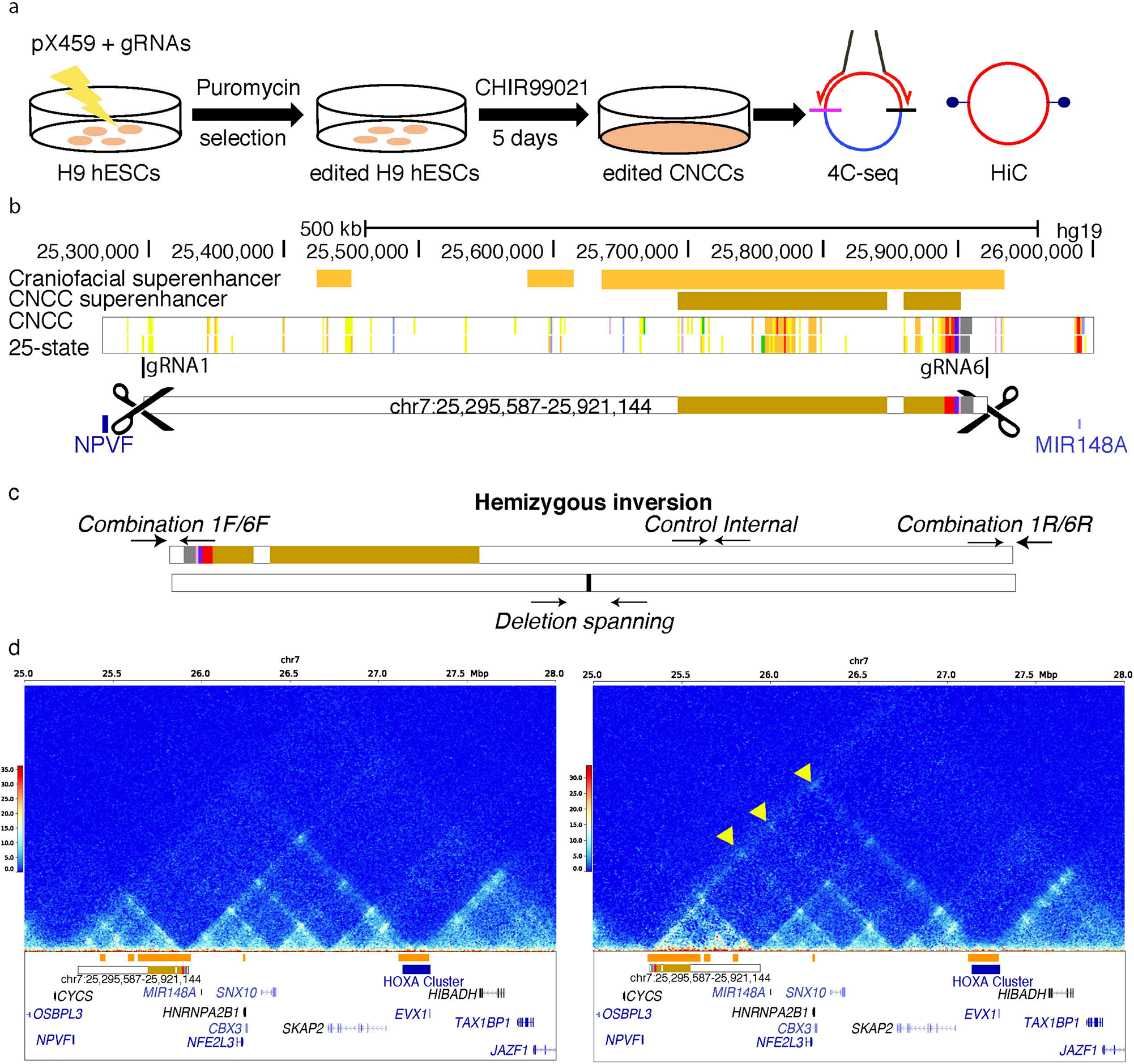
Editing of hESCs resulted in inversion of superenhancer target. Method of genome editing H9 cells (a). Location of guide RNAs gRNA1 and gRNA6 relative to the WT orientation (b). Screening strategy for determining whether clones are heterozygous for the 1/6 deletion and determining if a clone contains an inversion of the targeted region, orientation of hemizygous inversion clone is shown (c). HiC interactions in H9-derived CNCCs from WT (d-left panel) and clone with hemizygous inversion (d-right panel). The HiC plot made for INV used alignment to a custom version of the hg19 genome with the specific inversion on chromosome 7 introduced. New contacts created by inversion are marked with yellow arrows.

Interestingly, we did identify one clone which had lost one allele of the region and inverted the other, resulting in a hemizygous inversion illustrated in Figure 4c (also see **Figure 4 Supplement 2**). All clones we obtained grew relatively normally in ESC culture conditions and we did not notice any major difference in morphology or gene expression (**Figure 4 Supplement 3**). Given that HSPCs lacking a copy of the TAD boundary had altered differentiation characteristics relative to wild type controls (Zhang et al., 2020) we proceeded with experiments to derive CNCCs. When these ESCs were directed down the CNCC lineage, both the heterozygous cells and the cells harboring the single inverted allele robustly differentiated and were very similar to WT cells (**Supplemental Table 8**). When we examined gene expression, we found little difference in global gene expression or genes surrounding the locus (**Figure 4 Supplement 4; Supplemental Table 9**). This suggested that the *HOXA* gene cluster did not adopt new interactions in the absence of this region. However, the inversion cell line resulted in considerably higher contact frequencies between the *HOXA* gene cluster and the now more distant TAD boundary (**Figure 4d**.). This was particularly surprising as this TAD boundary had been moved nearly 600kb farther away from its target and the CTCF motif orientations had been inverted relative to the *HOXA* gene cluster. Given these results we reasoned that *HOXA* gene expression might be maintained at normal levels in these cell lines through upregulation of the cluster that remained in *cis* with the superenhancer region.

### Syntenic superenhancer cluster in mouse makes 3-dimensional contacts with *Hoxa* cluster

While these results have implications for models of chromatin architecture and loop formation, the robust expression of *HOXA* despite these changes precluded us from making any determination of the role of this region in craniofacial related biology. We therefore asked whether this region might have similar epigenomic properties in mice, and if it could be better studied in a system where we could generate homozygous null animals.

Syntenic regions often have preserved regulatory structure and features (Dixon et al., 2012; Vietri Rudan et al., 2015) and this was indeed the case for this craniofacial specific superenhancer. This region is part of a large syntenic block between the two species which stretches nearly 10 Mb in length with the *HOXA* gene cluster roughly at its center (**Figure 5 Supplement 1**). When we compared chromatin states using an 18 state chromHMM model on epigenomic data obtained by Mouse ENCODE across multiple tissues and stages of development (Methods) we found similar trends in activation across the larger gene desert region (**Figure 5 Supplement 2**). Chromatin state segmentation of mouse craniofacial tissue (E9.5-E15.5) demonstrated that the *Hoxa* cluster and the 2Mb adjacent to the anterior side have similar chromatin state characteristics to that of human embryonic craniofacial tissue and the validated craniofacial enhancers (**Figure 5, Figure 5 Supplement 3**). When we analyzed the mouse epigenomic craniofacial data from developmental stages comparable to our human tissue samples, similar portions of the gene desert between *Npvf* and *Nfe2l3* were identified as superenhancers (**Figure 5**).

**Figure 5.**
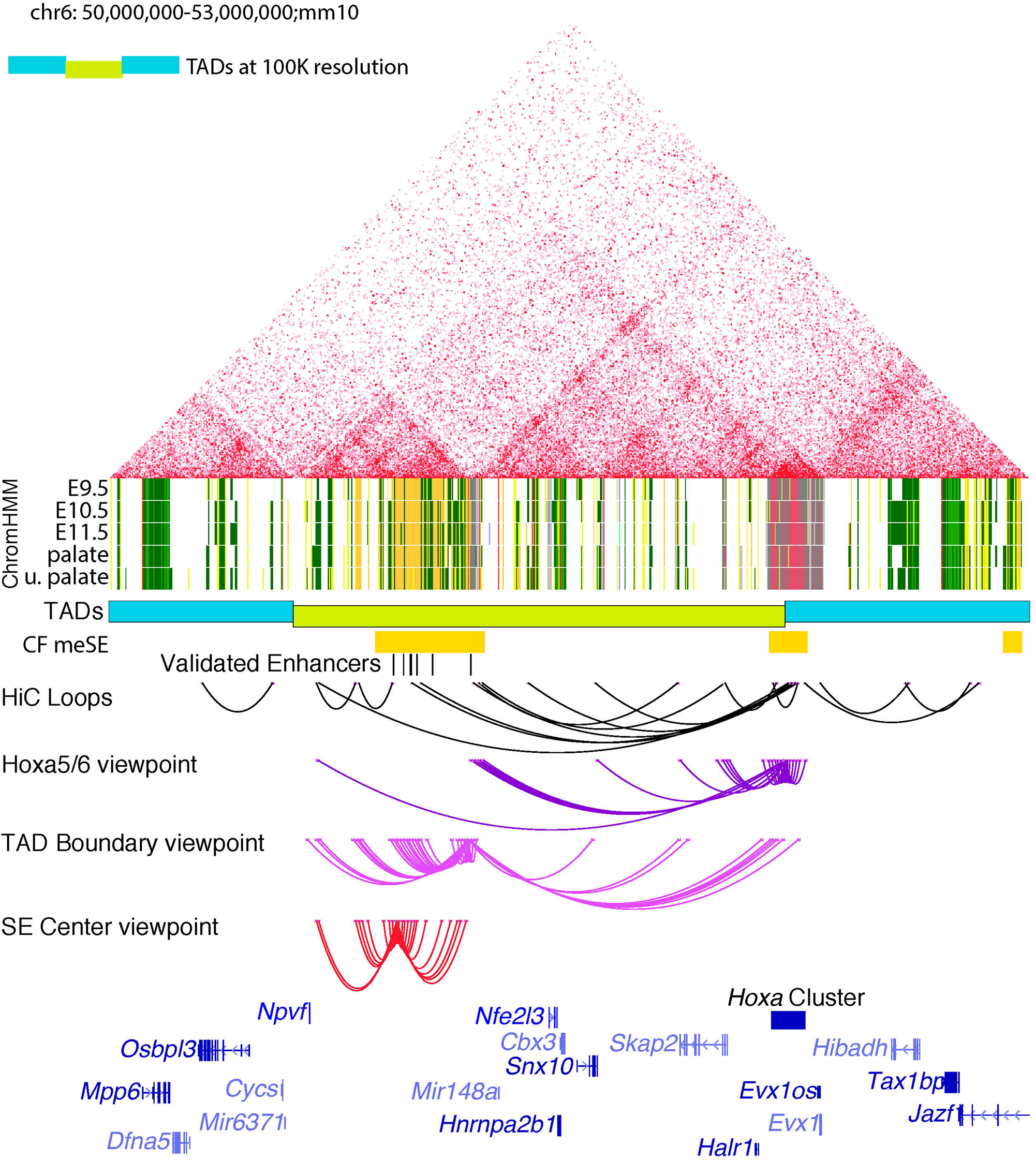
Chromatin Architecture in Primary Mouse Embryonic Craniofacial Tissue. HiC of E11.5 mouse embryonic craniofacial tissue. TADs called at 100Kb (blue/dark yellow). Mouse embryonic craniofacial superenhancers (CF meSE) determined by ROSE algorithm. Enhancer segments with validated craniofacial activity (shown in Figure 2), human enhancer segment coordinates arrived at via liftover. ChromHMM chromatin states for the 18-state model of embryonic craniofacial tissue for E9.5-E11.5, E12.5 palate and E13.5 upper palate; individual enhancer segments are yellow and orange. HiC loops as predicted at 10Kb resolution, 4C-seq loops at 10Kb resolution. Viewpoint near intergenic space between Hoxa5 and Hoxa6 (purple), viewpoint at TAD boundary (magenta), and viewpoint at center of superenhancer subTAD (red).

Having demonstrated conservation of chromatin states between human and mouse craniofacial development, we wondered if the three dimensional structure of the region is also conserved. Utilizing circularized chromosome conformation capture with sequencing (van de Werken et al., 2012) (4C-seq) we assessed the interactions of four viewpoints in this window in E11.5 mouse craniofacial tissue. For two viewpoints we identified extensive interactions within the identified window that do not cross the putative TAD boundary. When we assessed viewpoints flanking the TAD boundary, one of which contained the active enhancer HACNS50, we observed interactions within this identified region as well as significant interactions with the *Hoxa* gene cluster (**Figure 5)**. To confirm these interactions, we performed additional 4C-seq experiments utilizing viewpoints located directly within the *Hoxa* cluster and the promoter of the *Skap2* gene. We observed strong interactions between both of these viewpoints and the TAD boundary of the original window **(Figure 5 Supplement 3, lower panel**). Interestingly, *Hoxa* made contacts with the outer limits of this window but contacts were not observed within the window. These findings illustrate that the region we identified in human craniofacial tissue makes strong contacts over nearly 1.5 Mb with genes of the *Hoxa* cluster in developing mouse craniofacial tissue. Overall these findings indicate this region is conserved at multiple levels: primary sequence within the superenhancer region, genomic position relative to potential regulatory targets, chromatin activation in tissues across developmental stages, and long-range three dimensional contacts.

### Deletion of the *Hoxa* global control region (GCR)

Conservation of so many functional aspects pointed to this region being a global control region (GCR) for the *Hoxa* gene cluster and likely very important for craniofacial development. Thus we proceeded with making an orthologous deletion of this potential GCR in mouse to study the effects of this superenhancer region on craniofacial development. We identified guide RNAs in the mouse genome very close to the orthologous positions utilized in the human cells (**Figure 6a**; see Methods). These guides were then injected with Cas9 mRNA into fertilized mouse eggs. This resulted in one heterozygous founder which was genotyped by PCR and DNA sequencing, then bred to produce the F1 generation. This resulted in five heterozygous F1 mice (3 female, 2 male) used to establish our breeding colony. The F1 mice were overtly normal and fertile. Since this was such a large genetic manipulation and to eliminate interference between potential off target effects we performed three backcrosses utilizing C57B6 wild type mice and heterozygous GCR deletion (ΔGCR/+) mice. The original F1 mice remained healthy during this time and we obtained expected numbers of heterozygous offspring across all backcrossing. We then crossed heterozygous ΔGCR/+ mice and observed perinatal lethality in the first two days in all homozygous pups. These pups seemed grossly normal externally but when we examined the internal craniofacial structures we found 55% (n=6/11) with clefts of the secondary palate. (**Supplemental Table 10**). We inspected embryos at multiple stages of development and did not observe any distinct morphological differences before E14.5. However, at this stage the upper palate frequently failed to fuse in the ΔGCR/ΔGCR embryos. Limb staging and comparison of fetal weight at E17.5 did not indicate significant body-wide developmental differences in ΔGCR/ΔGCR mice compared to WT and ΔGCR/+ littermates (**Supplemental Table 10**).

**Figure 6.**
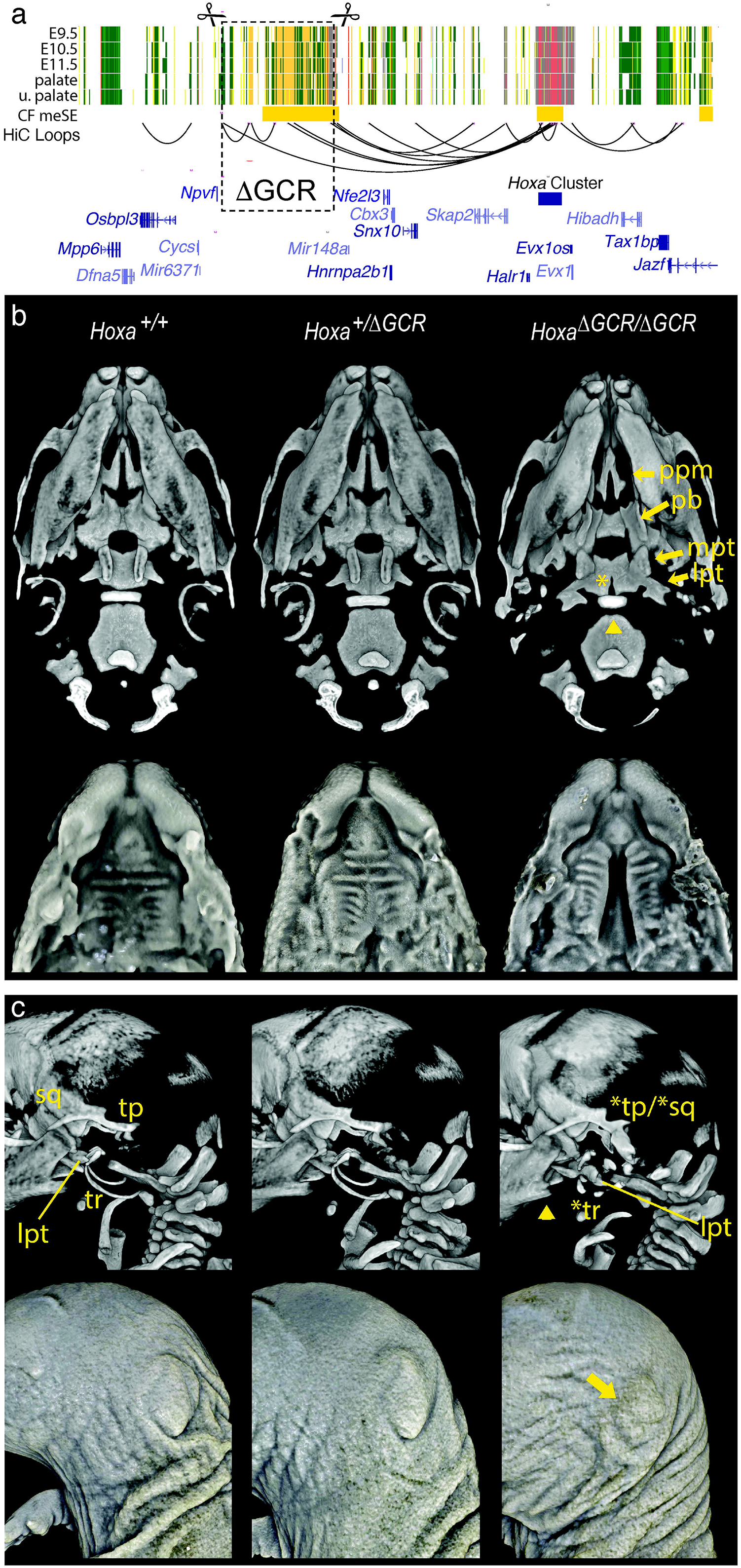
Deletion of the craniofacial-specific superenhancer distal to the Hoxa gene cluster mimics the Hoxa2 null phenotype. **A**. Schematic of deletion, mouse chr6:50673614-51196805 (mm10 or mm39?), spanning major predicted contacts with the *Hoxa* cluster. **B**. (upper row) Three-dimensional rendered images generated from microCT scans of wildtype E18.5 embryos and their heterozygote and homozygote ΔGCR littermates. Ventral view of the skulls reveals multiple cranial base and palatal bony defects in homozygotes. The palatal defects include cleft palate in ∼66% of homozygotes and reflected by marked separation of the palatine processes of the maxillae [ppm], separation and ventrally projecting palatine bones [pb], as well as lateral flattening of the medial pterygoid processes [mpt] of the basisphenoid. There was variability in the palatal presentation in homozygotes (see Figure 6 Supplement 1). The cranial base presentation, characterized by the notably abnormal appearance of the lateral pterygoids [lpt] of the basisphenoid and abnormal anterior shape of the basioccipital (denoted by the arrowhead), was fully penetrant in homozygotes. A posterior cleft (arrowhead) or small notch in the basisphenoid was also evident in ∼50% of homozygotes. **(lower row)** Soft tissue rendering from microCT scans of E17.5 embryos confirm the cleft palate observed in some homozygotes. Note the normal formation of rugae despite the cleft. **C**. Left lateral view of the bony (top) and soft tissue (bottom) rendering of microCT scans of littermates. Homozygotes show mirror duplications of the tympanic ring (tr; *tr), tympanic process and squamous bone (tp/sq; *tp/*sq) reminiscent of previously reported *Hoxa2* null mce. The abnormal lateral pterygoid (lpt) of the basisphenoid is evident from this view of homozygotes. Although not previously described in *Hoxa2* null mice, the mandibular angle was consistently hypoplastic (arrowhead) in homozygotes. On the soft tissue renderings, variable severity microtia can be clearly seen in homozygotes (arrow). Microtia ranges from grade I to grade III.

MicroCT scans of ΔGCR/ΔGCR embryos compared to WT and ΔGCR/+ littermates at E18.5 confirmed the incidence of cleft palate in homozygous embryos as well as striking anomalies of the cranial base and ears (**Supplemental Movie 1**). Notably, the ΔGCR/ΔGCR embryos also had hypoplastic mandibular angles, while the mandibular angle also appeared slightly smaller in ΔGCR/+ embryos compared to WT littermates. With regard to the cranial base, large bony appendages on the lateral pterygoid of the basisphenoid and lateral flattening of the medial pterygoid process were present in ΔGCR/ΔGCRembryos. (**Figure 6b upper, Supplemental Movie 2, Supplemental Movie 3**). In ΔGCR/ΔGCR embryos with cleft palate, ventral projections from the palatal bones instead of the normal horizontal projection toward the midline suggested that the palatal shelves had not elevated. However, soft tissue rendering from the same scans showed relatively normal rugae formation despite the failure of the shelves to approximate and the aberrant underlying palatal bone projections. The remaining ΔGCR/ΔGCR embryos without cleft palate exhibited fairly normal palatal bones, maxillary palatine processes and basal pterygoids (although one was mildly impacted) (**Figure 6b, lower**). Despite the variably penetrant clefting, all ΔGCR/ΔGCR embryos showed mirror duplication of the tympanic ring and potential partial mirror duplication of the tympanic process and squamosal bone (**Figure 6c upper**; **Supplemental Movie 4**). The external ear (pinna) was also overtly microtic in all ΔGCR/ΔGCR embryos, although the severity varied from embryo to embryo. The bony and soft tissue ear phenotypes did not correlate with palatal clefting (**Figure 6c, lower**). Collectively, the developmental phenotypes seen in the ΔGCR/ΔGCR mice were essentially identical to those reported in *Hoxa2*-/-lines (Rijli et al., 1993; Gendron-Maguire et al., 1993; Barrow and Capecchi, 1999) suggesting that the lack of the GCR had very specific gene expression consequences.

To determine if this was the case, we collected multiple tissues from embryos at E11.5. This stage is before the differences in phenotype are apparent and when *Hoxa* gene expression is robust (Donaldson et al., 2012). When we compared global gene expression of craniofacial tissue between ΔGCR/ΔGCR and wildtype type embryos we found highly specific effects on *Hoxa2, Hoxa3, Hoxaas2* (**Figure 7a**). We did not observe any significant changes in expression for other genes surrounding the deletion nor residing between the deletion and the *Hoxa* gene cluster (**Figure 7b**). When we examined heart and limb tissue we found a small number of differentially expressed genes between ΔGCR/ΔGCR and wild type mice, however none of these were at this locus and the *Hoxa* gene cluster maintained consistent expression across all embyros (**Figure 7 Supplement 1**). Inspection of the three-dimensional architecture of this region using HiC in craniofacial tissue from ΔGCR/ΔGCR mice showed that we were able to achieve a complete deletion of the TAD subdomain containing the GCR (**Figure 7 Supplement 2**). This resulted in a smaller TAD domain that excludes the anterior half of the *Hoxa* gene cluster but otherwise did not cause further enhancer adoption or large scale changes in chromosome conformation. These findings were consistent with the very specific effect on expression of the Hoxa gene cluster and none of the other nearby genes. Overall we have shown that deletion of an entirely noncoding craniofacial-specific superenhancer region has tissue-specific consequences on a small number of genes located a considerable distance away. Mice completely lacking this region frequently had craniofacial abnormalities and died soon after birth.

**Figure 7.**
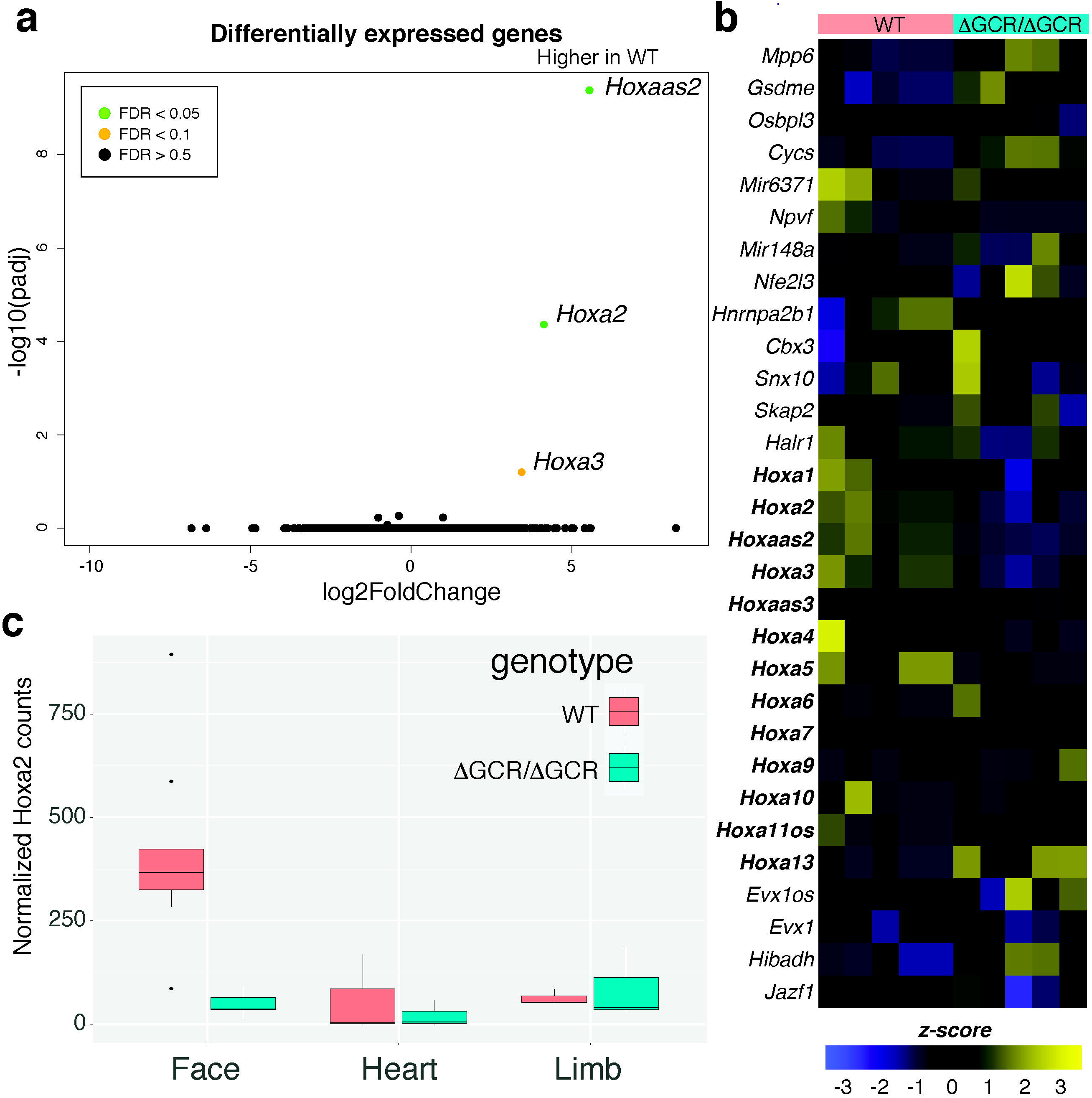
Deletion of a superenhancer region has specific effect on *Hoxa* genes. a. Deletion resulted in decrease in Hoxa gene expression without similar decrease in expression of intervening genes such as Snx10 and Skap2. b.Heatmap of expression from all replicates of WT and ΔGCR/ΔGCR for genes indicated in panel a. c. *Hoxa2* expression was substantially different in E11.5 ΔGCR/ΔGCR vs. WT littermates in craniofacial tissue but not in heart or limb.

### Identification of novel deletion of TAD boundary in a human fetus

Given the strong phenotype in the mouse homozygous knockout, we wondered if similar phenotypes might be apparent in human development. The analysis of the DECIPHER database described above indicated that indeed this region is associated with craniofacial abnormalities. However, all of the copy number changes were quite large (> 3MB) and included many genes outside of the noncoding region of interest. Thus it remained unclear if alteration of this region alone could result in craniofacial abnormalities. Recently, we identified a *de novo* deletion partially overlapping the *HOXA* superenhancer region in a fetus with severe craniofacial abnormalities (**Figure 8**). Following targeted ultrasonography (gynecologists report in Supplemental Text), the pregnancy was terminated at 14 weeks. The autopsy revealed bilateral cleft lip and palate, an underdeveloped nose with a single nostril, as well as clubfeet and anal atresia. Shallow whole genome sequencing identified a 550 kb deletion (chr7:25800001-26350000, hg38), which was absent in the parents and encompasses the *NFE2L3, HNRNPA2B1, CBX3, MIR148A* genes, as well as exons 1 and 2 of the *SNX10* gene (**Figure 8** and **Figure 8 Supplement 1**). Mendeliome analysis for this trio did not reveal any other putatively causal variants. As discussed above, none of the genes directly affected by the deletion seem appropriate candidates to explain the severe craniofacial defects we observed in this fetus. Interestingly, the deletion overlaps the 3’ end of the *HOXA* superenhancer we identified, including the 3’ TAD boundary, which forms strong interactions with the *HOXA* gene cluster, and the validated HACNS50 enhancer element, which is active in mouse embryonic craniofacial and limb tissue. The role of *HOXA* genes during craniofacial development, the craniofacial abnormalities occurring in *HOXA*-associated syndromes, the tissue-specificity of the affected superenhancer region and its strong interaction with the *HOXA* cluster, as well as the association of CNVs affecting the superenhancer with craniofacial malformations, all implicate *HOXA* dysregulation as the underlying cause of the severe craniofacial defects in this fetus. However, further experiments are needed to conclusively rule out that the partial deletion of the *SNX10* gene, which has been associated with autosomal recessive osteopetrosis and also includes clinical features such as macrocephaly and facial anomalies (OMIM #615085), contributes to the phenotype.

**Figure 8.**
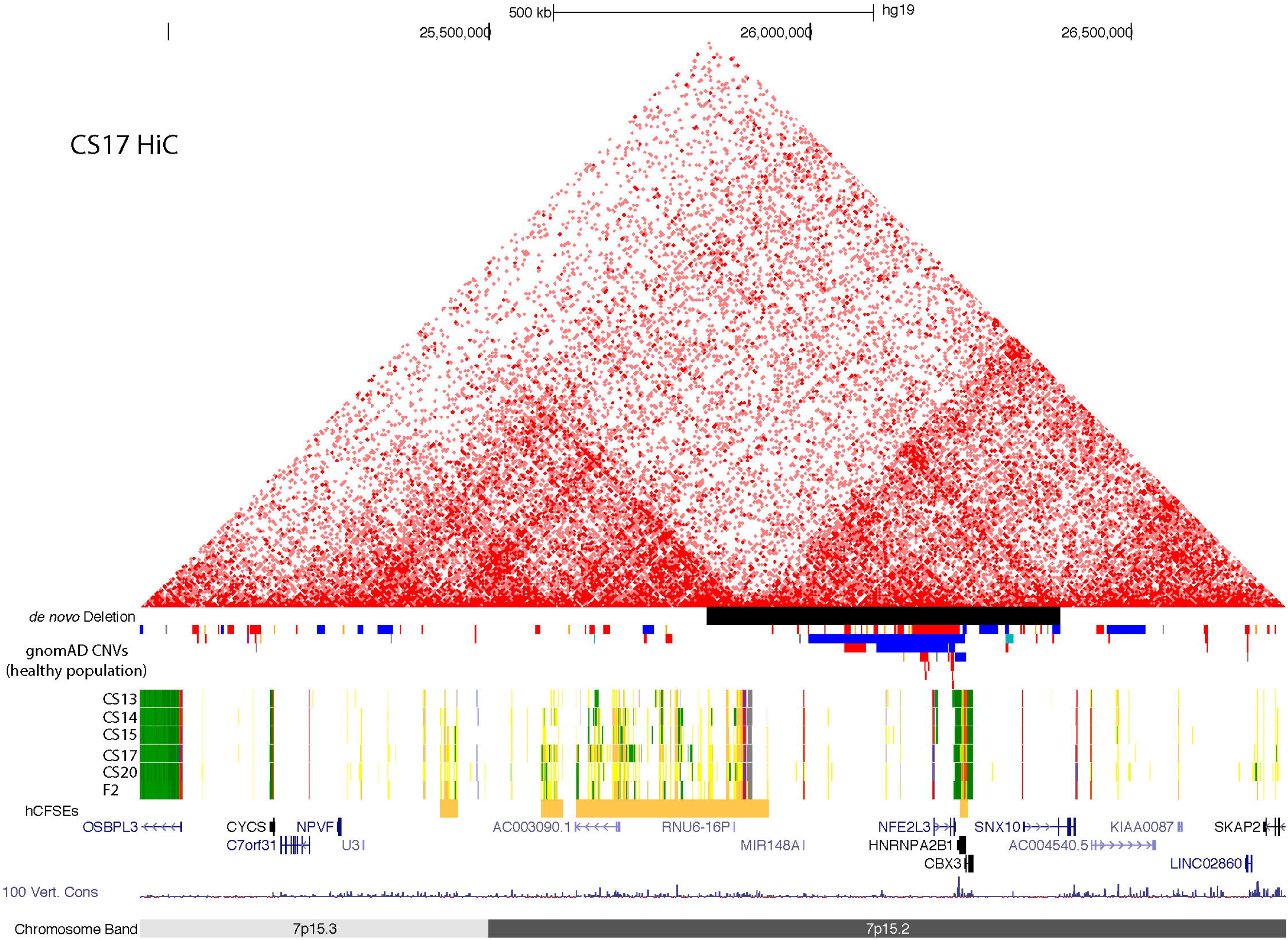
Location of de novo deletion overlapping GCR. Browser image (hg19) showing de novo deletion presented by the group at Ghent University corresponding to Chr7:25,839,621-26,389,620 on the hg19 assembly. Also shown are the UCSC Browser tracks for the 25-state model, human embryonic craniofacial superenhancers and gnomAD Structural Variation track filtered to show CNVs >300bp. Colors in the gnomAD track are as they appear in the UCSC genome browser, red bars signify deletions and blue bars duplications.

## Discussion

Collections of coactivated enhancers commonly referred to as superenhancers encompass genes that play important roles in tissue and disease specific biology. Our identification and global analysis of superenhancers active in craniofacial development largely confirmed these findings, and demonstrated that many craniofacial related transcription factors are likely controlled by such regions. Moreover, we found that superenhancers specific to craniofacial development are enriched for transcription factors and other genes that are directly linked to craniofacial related diseases. These included *ALX4, MSX2*, and CYP26 family members, which demonstrated clear craniofacial-specific regulatory landscapes. However many craniofacial-specific superenhancers we identified neither included nor were in close proximity to known craniofacial genes.

We reasoned that such regions might either contain a previously unannotated gene or operate over long distances to target a craniofacial relevant gene. Unlike individual enhancers, the regions we identified occupy fairly large portions of the genome, making them challenging to study as a complete unit. Furthermore, deletion of many individual enhancers and even megabase-scale, completely noncoding regions have resulted in mice with no overt phenotype (Nóbrega et al., 2004; Ahituv et al., 2007; Attanasio et al., 2013; Dave et al., 2017; Moorthy et al., 2017). These findings call into question the role such regions might play during development. However, we observed functional gene enrichments in the surrounding areas that suggest an important role for these regions in craniofacial development and disease.

To address this we focused on completely noncoding superenhancers whose removal was not predicted to disrupt a gene. We prioritized regions by overlaying multiple layers of functional genomics data and publicly available enhancer validation experiments and tested one of the highest priority regions. This region had nearly one hundred active craniofacial enhancer segments, many VISTA validated craniofacial enhancers, showed functional conservation in development between both human and mouse, and was embedded in a large syntenic region that also included the *HOXA* gene cluster. Examination of chromosome conformation data revealed that this region forms a distinct topologically associated domain (TAD) in most tissues. Interestingly, in craniofacial tissue this region becomes a sub-TAD of a larger domain that includes the HOXA gene cluster.

TAD organization has been shown to be important for gene regulation. Breaking or inverting TAD boundaries to affect their enhancer-gene content can result in developmental abnormalities (Lupiáñez et al., 2015; Melo et al., 2020, 2021; Symmons et al., 2016). In an attempt to separate the function of the novel superenhancer region from larger scale TAD organization, we attempted to create a “scarless” edit with respect to TAD boundaries. We targeted strong CTCF binding sites at both TAD boundaries in human cells. While we were unable to create homozygous deletions in human embryonic stem cells, we identified a cell line that lacked one copy of the region and harbored a ∼600kb inversion of the remaining copy. This inversion showed that this region functions as an enhancer in the classical sense, functioning to regulate gene expression over long distances independent of orientation. NCC differentiation of this inversion line did not reveal significant differences in gene expression of the *HOXA* gene cluster, nor any other genes in the direct vicinity.

Interestingly, we observed an increased frequency of interactions between the TAD boundary that was now 600kb further away from the HOXA gene cluster and the anterior *HOXA* genes. This suggests that this boundary in particular is programmed to target the *HOXA* gene cluster in neural crest and can function over even greater distances, even dominating over other boundary regions that are closer to the target and have properly oriented CTCF motifs for looping (Dixon et al., 2012; Guo et al., 2015; Nanni et al., 2020). While the lack of impact on gene expression was unexpected, our inability to identify homozygous knock-outs in H9 ESCs and the behavior of the inversion suggested that the absence of this region might have a strong impact on *HOXA* gene expression and is essential for human stem cell survival. Recent work in hematopoietic stem and progenitor cells (HSPCs) support this conclusion (Zhang et al., 2020), where a 17kb deletion encompassing the strong TAD boundary more proximal to the *HOXA* gene cluster led to an increase in differentiated cells. This suggests a role for the interaction between the anchor and the *HOXA* cluster in the maintenance of stem cell identity. Disruption of this region, identified in HSPCs as a DNA methylation canyon, led to altered expression of *HOXA* genes without significant changes in expression for the intervening genes. These results along with our findings point to this region being a potential global control region of the *HOXA* gene cluster, similar to those already known to control other HOX gene clusters (Spitz et al., 2003) (Kleinjan and Lettice, 2008; Lee et al., 2006).

Our experiments in cell culture, while suggestive, failed to address the original question of whether specific superenhancers are important for development. We therefore generated an orthologous deletion in mouse. While the heterozygous founders and many resulting progeny lacked any clear phenotype, homozygous mice never survived postnatally beyond 48 hours and frequently had clefts of the secondary palate. Gene expression analysis of craniofacial tissues at E11.5, prior to the emergence of clefting phenotypes, showed specific effects on the anterior *Hoxa* genes, particularly *Hoxa2*, and no other genes in the region. Other tissues from the same stage of development showed no such issue in gene expression. Given the strong effects on *Hoxa2* gene expression in our mice and the documented role of *HOXA2* in craniofacial development, we more closely examined the skeleton of homozygous deletion mice using microCT. We found craniofacial bony phenotypes including malformed basisphenoid bone, tympanic process and tympanic ring defects, as well as microtia in the Hoxa GCR null mice we generated. These were in striking similarity to those seen in previously published *Hoxa2* null mice (Rijli et al., 1993; Gendron-Maguire et al., 1993; Barrow and Capecchi, 1999; Santagati et al., 2005).

These results in mice along with the human fetus we described above point to this superenhancer region being important for craniofacial *HOXA* gene expression and essential for life in both humans and mice. Within the region of the human deletion, the only genes with comparable constraint to anterior *HOXA* genes are *CBX3* and *HNRNPA2B1* (LOEUF score 0.67, 0.22 respectively). However, haploinsufficiency of either gene does not produce phenotypes similar to the described fetus (Kim et al., 2013; Aydin et al., 2015) (Qi et al., 2017; Gillentine et al., 2021). The other distinct difference between human and mice is that loss of a single allele of the GCR in mice does not result in any overt phenotype, whereas perturbing a single allele in humans can lead to catastrophic effects. As we noted above, a sequence with a high number of human-specific substitutions, HACNS50, is located in close proximity to the TAD boundary of the GCR (Prabhakar et al., 2008). This could indicate that this region has gained or lost activities specifically in humans, resulting in human-specific sensitivity to loss of this region.

Further dissection of the *HOXA* GCR is required to determine whether there are one or a few enhancer elements primarily responsible for the regulatory activity and lethality. Given the specific distribution and functions of the anterior *HOXA* genes, it will be important to determine whether portions of the superenhancer have distinct control over expression of specific *HOXA* genes. Deletion of select elements could impact the limbs or other organs despite not being affected by deletion of the entire superenhancer. CRISPR-Cas9 directed dissection of the superenhancer cluster coupled with single-cell ATAC-seq provides a novel direction to identify whether there are developmental- and cell type-specific modules within the HOXA GCR.

## Acknowledgments

We would like to thank the laboratory of James P. Noonan for providing images of the HACNS50 enhancer reporter experiments. We would also like to thank the staff of the University of Connecticut Computational Biology Core and the High Performance Computing Center for assistance with package installation and software/hardware support. This work was funded by the National Institutes of Health NIDCR 1R01DE028945 (JC), NIDCR 1R03DE028588 (JC), and NIGMS 5R35GM119465 (JC). This study makes use of data generated by the DECIPHER community. A full list of centres who contributed to the generation of the data is available from https://deciphergenomics.org/about/stats and via email from. Funding for the DECIPHER project was provided by Wellcome.

## Author Contributions

Conceptualization AW and JC. Investigation AW, ED, MB, TNY, EWW, and TC. Formal Analysis AW, ED, MB, TC, SG, SV, and JC. Resources NG, ER, JVD. Writing - Original Draft AW and JC. Writing - Review and Editing AW, ED, MB, TNY, EWW, ER, JVD, SV, TC and JC. Supervision SV, TC and JC. Funding Acquisition JC.

## Data Availability

Signal tracks for these experiments are available through track hub functionality at the UCSC Genome Browser (https://cotneylab.cam.uchc.edu/~jcotney/CRANIOFACIAL_HUB/Craniofacial_Data_Hub.txt), as a public session (https://genome.ucsc.edu/cgi-bin/hgTracks?hgS_doOtherUser=submit&hgS_otherUserName=Jcotney&hgS_otherUserSessionName=Human%20HOXA%20LCR) or at the Cotney Lab website (https://cotney.research.uchc.edu/hoxa_gcr/).

## Methods

### ROSE algorithm

We used the Rank Ordering of Super-Enhancers (ROSE) algorithm (Whyte et al., 2013; Lovén et al., 2013). Briefly, H3K27ac ChIP-seq and Input (control) data and genomic regions classified with active chromatin states were provided to the algorithm. For the 25-state model, these states included 1_TssA, 2_PromU. 3_PromD1, 4_PromD2, 9_TxReg, 10_TxEnh5’, 11_TxEnh3’, 12_TxEnhW, 13_EnhA1, 14_EnhA2, 15_EnhAF, 16_EnhW1, 17_EnhW2, 18_EnhAc, 19_DNase, 22_PromP, and 23_PromBiv. For the 18-state model, these states included 1_TssA, 2_TssFlnk, 3_TssFlnkU, 4_TssFlnkD, 7_EnhG1, 8_EnhG2, 9_EnhA1, 10_EnhA2, 11_EnhWk, 14_TssBiv and 15_EnhBiv. Enhancers within 12.5kb were “stitched” together as a single region. “Stitched” enhancers as well as enhancers without another enhancer within 12.5kb were ranked for H3K27ac enrichment. Determination of superenhancer was based on the threshold of where the slope of H3K27ac enrichment = 1.

### Gene Ontology and Disease Ontology

Biological Process Gene Ontology and Disease Ontology from DisGeNet enrichment was primarily performed with the R packages clusterProfiler (3.14.3) and DOSE (3.12.0) (Yu et al., 2012). Genes overlapping TSSs or assignment of nearest genes was performed using BEDTools (2.29.0) (Quinlan and Hall, 2010). LiftOver analysis was performed using KentUtils (1.04.00). In some cases Disease Ontology enrichment was identified through WEB-based GEne SeT AnaLysis Toolkit (WebGestalt) (Zhang et al., 2005; Liao et al., 2019; webgestalt.org).

### In-vivo validation of HACNS50

In vitro lacZ reporter assay of an additional enhancer element was performed by Justin Cotney. A 2.6 kb segment centered on the conserved sequence corresponding to HACNS50 (Prabhakar et al., 2008) was amplified from human genomic DNA by polymerase chain reaction (PCR) using the following primers: HACNS50 F 5’-CACCCCATTTCTGAGGGGGAAATAA-3’, HACNS50 R 5’-TTATTTCCTTCAGGCCCTTG-3’, and cloned into an Hsp68-lacZ reporter vector as previously described (Visel et al., 2008). Generation of transgenic mice at the Yale University Transgenic Mouse Facility and embryo staining were carried out as previously described (Visel et al., 2008). We required reporter gene expression in a given structure to be present in at least three independent transgenic embryos as assessed by two researchers to be considered reproducible.

### H9 hESC cell culture

Routine culture of H9 cells was done in feeder-free conditions using Matrigel substrate (Corning, 354277) and Essential 8 media (Gibco, A1517001). Where feeder cells were used (following nucleofection with gRNAs) the cells were plated on DR4 MEFs in a gelatinized 10cm tissue culture dish with hESC on MEF media (DMEM high glucose + 10% FBS).

### Guide RNA design and preparation

We used the online tool at http://crispor.tefor.net/ to select appropriate gRNAs. Guide RNA oligonucleotide pairs were synthesized by IDT (Coralville, IA, USA) and received in lyophilized form. The gRNA oligos were diluted to 100mM with sterile water. The oligonucleotide pairs for each gRNA were phosphorylated and annealed in a reaction containing 100mM of each oligonucleotide, 5U T4 PNK (NEB, M0201) and T4 ligation buffer at 37°C for 30 minutes with boiling at 95°C for 5 minutes followed by a gradual cool to 25°C at an approximate rate of 5°C per minute. The resulting annealed oligos were diluted 1:200 and added to a ligation reaction with 50ng BbsI digested pX459 plasmid, 1x Quick Ligation buffer and Quick Ligase (NEB, M2200). The ligation reaction proceeded for 10 minutes at room temperature and then was treated with PlasmidSafe Exonuclease. The Exonuclease reaction contained the full volume of the ligation reaction, 1X PlasmidSafe Buffer, 1mM ATP, and 3.2U PlasmidSafe Exonuclease (Epicentre, E3101K) and was incubated at 37°C for 30 minutes. The recombinant plasmids were introduced into competent dh5alpha cells via heat shock at 42°C, recovered in SOC at 37°C for 60 minutes with gentle rocking and plated on LB-Ampicillin plates. After overnight incubation of the plates at 37°C in a bacterial incubator, colonies were selected and grown overnight in 3ml liquid LB-Ampicillin at 37°C in a shaking incubator. Glycerol stocks were prepared from a fraction of each culture. Plasmid DNA was extracted from the overnight cultures using Qiagen miniprep kit (Qiagen, 27106) according to the manufacturer’s instructions for centrifuge column prep. Quantity of DNA was measured using Nanodrop spectrophotometer and proper construction of plasmid was confirmed using an EcoRI/BbsI double digest. Plasmid preps with the proper restriction digest profile were sent to Genewiz (South Plainfield, NJ, USA) for sequence confirmation by Sanger sequencing. The plasmid DNA was sent pre-mixed with LKO.1 5’_U6primer (5’-GACTATCATA TGCTTACCGT-3’). Glycerol stocks of the plasmids that were confirmed correct by Sanger sequencing were grown and prepared for maxiprep plasmid isolation (Zyppy Plasmid Maxiprep Kit, Zymo 6431). Proper cloning was achieved for all gRNAs.

### Genome editing of H9 hESCs

Guide RNAs gRNA1 and gRNA6, which together would cleave out the region chr7:25,295,587-25,921,144 (hg19) were introduced into H9 hESCs by nucleofection (Lonza, Basel, Switzerland). To start, hESCs in one confluent well of a 6-well plate were singlized, first detached from the Matrigel plate with Accutase (Thermo Fisher, 00-4555-56) at 37°C for 10 minutes, and pipetted to single cell suspension in Essential 8 media + 10mM ROCK inhibitor (Tocris Bioscience, Y-27632 dihydrochloride). Approximately half of the cells were transferred to a 15ml conical tube and centrifuged at 1000rpm for 5 minutes. The media was completely removed and replaced by 100ml of Nucleofection Mix (82ml Nucelofector solution, 18ml supplement, 2.5mg gRNA1, 2.5mg gRNA6. The cells were resuspended in the Nucleofection Mix with a p200 pipette tip by pipetting gently three times. The sample was transferred to the cuvette using the 2ml pipette provided with the Nucleofection kit. The cells were placed in the Nucleofector and run on the program for hESC, P3 primary cell protocol. Following the time in the machine, 500ml hESC on MEF media plus ROCK inhibitor was added to the cuvette. The cells were plated on 10cm plates with Matrigel or DR4 MEFs. The cells were fed with the appropriate media plus ROCK inhibitor until colonies were visible. Selection with puromycin and support with ROCK inhibitor began the day following Nucleofection and continued for a second day. Subsequently only ROCK inhibitor was added to the media until colonies were visible. Colonies were screened via the hotshot method. Briefly, portions of colonies were picked and transferred with less than 200ml media to a PCR tube within a strip of 8 tubes. The samples were centrifuged for 2 minutes or longer in a microfuge with PCR strip tube adapter. The media was carefully aspirated and the cells resuspended in 20ml lysis buffer (25mM NaOH, 200mM EDTA). The cells were incubated for 45 minutes at 96°C in a thermalcycler. Following the lysis, freshly made neutralization buffer (40mM Tris, pH 5.0) was added to each tube and 5ml of DNA was used as a template for the screening PCR reaction using primers. Clones identified as edited had the target ends amplified by PCR with high-fidelity Taq polymerase (Pfusion HF; NEB, M030) and sent for Sanger sequencing (Genewiz, South Plainfield, NJ, USA). Sequences of gRNAs and screening primers are in **Supplemental Table 11**.

### Neural crest differentiation

Starting with one confluent well of a 6-well plate of H9 hESCs on Matrigel, the cells were detached using Accutase at 37C for 10 minutes and the cells pipetted into a single cell suspension using a 2ml serological pipette and resuspended into a total volume of 10ml of Essential 8 media. The cells were then pelleted by centrifugation for 3 minutes at 1000xg. The supernatant was aspirated and replaced with NCC Media (DMEM/F12 (Gibco, 10565-018) plus B27 supplement (Gibco, 17504-044 and Penicillin/streptomycin (Life Technologies, 15140122) with 3uM Chiron (CHIR-99021; Tocris Bioscience) and 10uM ROCK inhibitor. The cells were resuspended by gentle pipetting with a serological pipette and passed through a 40micron filter. The cells were counted with a hemocytometer and diluted to the desired density (30,000 cells per cm^2^) using the NCC Media with Chiron and ROCK inhibitor. Media was changed daily with NCC Media plus Chiron and ROCK inhibitor on the day following plating and NCC Media plus Chiron only for the remaining days. Differentiation is complete by day 5.

### Human primary tissue source

Pharyngeal arch and craniofacial tissue from unfixed, unsectioned embryonic tissue was obtained from the Human Developmental Biology Resource (HDBR), a part of the Wellcome Trust, in the UK.

### Human primary tissue fixation for HiC

Prior to beginning, all appropriate precautions were taken for the handling of potentially hazardous biological material. Individuals handling the samples wore disposable isolation gowns, disposable hair nets, disposable plastic face shields, disposable shoe covers and double layers of examination gloves. Surfaces and instruments involved in the processing of the samples were disinfected with 10% bleach afterwards. Samples were removed from -80 storage and removed from the tube by thawing in a small amount of PBS. The tissue was transferred to a dish containing cold PBS on a chilled microscope stage. The tissue was documented by photography through the microscope at several angles and photos of the tube were taken as well. The tissue was assessed as to whether it needed to be further dissected to isolate the specific tissue of interest and only the tissue of interest was transferred to a tube with 1mL PBS. The tissue was homogenized with an electronic tissue disruptor (Polytron PT 1200E, Kinematica, Fisher Scientific, USA). For HiC, formaldehyde was added to the remaining volume of homogenized tissue to a final concentration of 1%, incubated at room temperature for 15 minutes with rotation, quenched with 1.5M glycine added to a final concentration of 150mM and incubated for 10 minutes at room temperature with rotation. The cells were pelleted by centrifugation at 2500xg for 5 minutes at 4C. The supernatant was removed and discarded as formaldehyde waste and washed once with 1mL cold PBS. The cells were pelleted again by centrifugation as before, the supernatant removed and the fixed pellet frozen in a dry ice/ethanol slurry. The fixed samples and samples in Qiazol were stored at -80C until time of use.

### HiC

Crosslinked cell pellets or crosslinked embryonic tissue was resuspended in 1x Cell lysis buffer as used for 4C and nuclei released in a prechilled dounce homogenizer by 10 strokes with a loose glass pestle, 20 minutes rest on ice followed by 40 strokes with a tight glass pestle as for 4C. The nuclei were pelleted by centrifugation at 2500xg at 4C for 5 minutes, washed twice with 1x NEBuffer 3.1 and pelleted again as before. The nuclei were resuspended in 1x NEBuffer 3.1 and permeabilized by the addition of SDS to 0.1% SDS final concentration, incubated for 10 minutes at 65C with shaking at 800rpm on a ThermoMixer. The SDS was neutralized with Triton X-100 to 1% final concentration and incubated at 37C for 10 minutes with gentle rocking. Additional 10x NEBuffer 3.1 was added to adjust for the addition of SDS and Triton X-100 to restore the concentration to 1X NEBuffer 3.1 before adding 400U DpnII and incubated overnight at 37C with gentle rocking to digest chromatin. The next morning, DpnII was inactivated by incubation at 65C for 20 minutes. A quality check for the completeness of digestion was carried out using 10ul aliquots taken prior to the addition of DpnII on the previous day and following DpnII heat inactivation. The quality check aliquots were treated with 100ug Proteinase K for 30 minutes at 65C and liberated DNA extracted by phenol:chloroform extraction. The DNA was treated with 1ug RNAse A for 15 minutes at 37°C and then mixed with 6x loading dye and run on a 0.8% Agarose/0.5x TAE gel. If digestion was deemed complete, the remainder of the digested chromatin was prepared for ligation. The ends of the DNA were marked with biotin using a mix of non-biotinylated dCTP, dGTP, dTTP (250 uM final concentration of each) and 250 uM final concentration of biotin-14-dATP and 50U DNA polymerase I Large (Klenow) Fragment (NEB, M0210) within 1x NEBuffer 3.1, incubated at 23C for 4 hours in a ThermoMixer (900 rpm mixing; 15 secs every 5 mins). The biotinylated DNA was ligated in a reaction mixture containing 1x ligation buffer (NEB, B0202), a final concentration of 1% Triton X-100, a final concentration of 120ug BSA and 4,000U T4 DNA ligase (NEB, M0202) at 16C for 4 hours with gentle rocking. Following ligation, the samples were incubated with 500ug Proteinase K overnight at 65C. The following day an additional 500ug Proteinase K was added to each tube and incubated at 65C for 2 hours to ensure complete digestion of proteins and liberation of ligated DNA. The DNA was isolated by phenol:chloroform extraction using 15ml PhaseLock tubes and precipitated using 1/10 volume of 3M sodium acetate, pH 5.2 and 2.5 volumes of cold 100% ethanol. The samples were incubated either on dry ice for 20 minutes or at -80C for 45minutes-1 hour until the liquid became viscous but not solid. The DNA was pelleted by centrifugation at 15,000xg for 30 minutes at 4C in a chilled fixed-angle rotor. The supernatant was decanted and the pellet allowed to air dry very briefly before resuspending in 1x TLE (10mM Tris-HCl, pH 8.0, 0.1mM EDTA, pH 8.0) and concentrated in an Amicon 30kDa cutoff centrifugal filters (Millipore, UFC503024). The column was centrifuged at 14,000xg in a tabletop centrifuge for 5 minutes at room temperature, then the column washed at least twice by the addition of 1x TLE and centrifugation again as before except at the last wash, when spun for 10 minutes. Following the last wash, the volume remaining in the column was adjusted to 100ul with 1x TLE and the column inverted in a new collection tube and spun at 1000xg for 2 minutes to collect the concentrated DNA. The recovered DNA was treated with 1ug RNAse A for 30 minutes at 37C and the DNA quantified by Qubit (Thermo Scientific, Q32850). An aliquot of 800ng of samples was set aside for quality control assay to check completeness of ligation. The quality control assay is a PCR followed by digestion with MboI, ClaI or a double digest as described in the protocol by the Dekker lab (Belaghzal et al., 2017) and specific for DpnII digested chromatin. With extent of successful ligation confirmed, the biotin was removed from unligated ends in the reaction by incubation of 5ug of the ligated DNA with 25uM each of dATP and dGTP (non-biotinylated) and 15 Units of T4 DNA polymerase in 1x NEBuffer 2.1 and incubated in a thermalcycler set to 20C for 4 hours. The enzyme was inactivated at 75C for 20 minutes and the sample cooled down to 4C. Multiple biotin removal reactions (up to 8) were set up and pooled afterwards and the volume adjusted to 500ul with molecular biology-grade water. The DNA was cleaned and concentrated in an Amicon column by centrifugation at 14000xg for 5 minutes and washed twice with 400ul molecular biology-grade water and centrifugation at 14000xg for 5 minutes. The DNA was recovered by inverting the column into a clean collection tube and centrifuging at 1000xg for 2 minutes. The volume was adjusted to 132ul after recovery using molecular biology-grade water and 2ul removed for sonication quality control as a pre-sonication check. The remaining 130ul of recovered DNA was transferred to a nonstick 1.5ml tube (Ambion, AM12475) and sonicated in a QSonica instrument (model Q-800R1-110) at 2C, amplitude 20% with 10 second pulses and 10 seconds of rest for 6 minutes at 10W per sample. Following sonication 2ul was removed as the post-sonication quality control sample. The results of sonication were checked using an Agilent Genomic DNA screentape. If the sheared DNA fell predominantly within the expected range of 200-400bp the samples could be further size selected using a two-step purification with Ampure XP beads to recover DNA fragments between 100 to 400bp. The DNA was enriched for fragments containing a ligation junction by capturing biotinylated fragments on MyOne Streptavadin C1 magnetic beads (Thermo Scientific, 65001). The DNA, still captured on the streptavidin beads, was used in the NEBNext UltraII End Prep reaction with the End Prep Enzyme Mix (NEB, E6050L) and incubated for 30 minutes at 20C followed by 30 minutes at 65C. Following the End Prep reaction, appropriate NEBNext Adaptors for Illumina (NEB, E7350) at 1:15 dilution were ligated using the NEBNext Ultra II Ligation Master Mix and incubated at 20C for 15 minutes in a thermalcycler without a heated lid to maintain proper temperature. The USER Enzyme was added to the ligation reaction and incubated at 37C for 15 minutes in a thermalcycler with the lid temperature at 50C. The beads were immobilized against a magnetic bar (Dynamag-2, Life Technologies, model # 12321D) and the beads washed twice with Tween wash buffer according to the MyOne Streptavadin C1 protocol, resuspended in 1x BB, made according to the MyOne Streptavadin C1 protocol, and washed twice with 1x TLE, then resuspended in 20ul TLE. The number of cycles required for the indexing reaction to amplify indexed, biotin-free DNA without introducing PCR bias was determined by setting up a small reaction using 3ul of beads with bound Adaptor Ligated DNA fragments in a 25ul indexing PCR reaction using the NEBNext Ultra II Q5 Master Mix, an Index Primer (i7 Primer), Universal PCR primer and 0.225ul 100x SybrGreen Dye (Invitrogen, S7563). The test PCR reaction was run in a (type of machine, brand, part number). The number of cycles to use in the indexing reaction for the remaining DNA-bound beads was determined by the number of cycles that gives 1/3 the maximum fluorescence. After determining the number of cycles, five 25ul indexing reactions were set up for each sample and the PCR run in a BioRad T100 Thermalcycler with the number of cycles adjusted. The completed indexing PCR reactions for a sample were pooled together in a 1.5ml non-stick tube and the streptdavidin beads immobilized on the Dynamag-2 magnetic bar. The supernatant containing the amplified indexed DNA was transferred to a new 1.5ml non-stick tube. A 3ul aliquot of the amplified DNA was saved as quality control sample for the removal of primer dimers. The rest of the amplified DNA was purified with Ampure XP beads using 1:1.5 ratio of sample to beads and eluted in 35ul TLE. The pre-purified quality control sample was compared to the post-purified sample using the Agilent Bioanalyzer D1000 Screentape. The molarity of the indexed library was calculated based on the NEBNext qPCR library quantification kit. Libraries were diluted to 4nM, denatured and prepared for sequencing on the NextSeq 500 or 550 with settings for single-index, paired-end sequencing with 36-42 cycles per end. Data was initially processed using HiC-Pro v.2.10.0 (Servant et al., 2015) visualization and prediction of TADs and loops were done using HiCExplorer v.3.7 (Ramírez et al., 2018).

### RNA extraction

Extraction of RNA from flash frozen cell pellets or primary tissue was carried out with the miRNeasy kit (Qiagen, 217004). The work surface, pipettes and centrifuge rotors were treated with RNAse Away (Life Technologies, 10328011) prior to beginning. Aliquots of reconstituted DNaseI were prepared from the RNase-free DNase Set (Qiagen, 79254) ahead of time and the appropriate amount of DNaseI in RDD buffer was prepared just prior to starting the extraction. Pellets or tissue were placed on ice and allowed to warm slightly, but not completely thaw. An average pellet or piece of primary embryonic tissue required 700ul Qiazol. Samples were homogenized in 700ul QIAzol by pipetting, brief vortexing and applied to Qiashredder columns (Qiagen, 79654). Homogenates were processed immediately after being allowed to incubate at room temperature for 5 minutes. The extraction proceeded with the addition of 140ul chloroform to the homogenate, which was then shaken vigorously for 15 seconds and allowed to rest at room temperature for 2-3 minutes. The homogenates were then centrifuged for 15 minutes at 12,000xg at 4C. The aqueous phase was transferred to a fresh 1.5ml tube (typically able to recover 300-350ul of aqueous phase). To the transferred aqueous phase 1.5 volumes of 100% ethanol was added and mixed by pipetting, then immediately passed through the RNeasy spin column and processed according to the manufacturer’s instructions with on-column DNase treatment and the addition of a second wash with Buffer RPE.

### RNA-seq library preparation

Total RNA quality was assessed using the Agilent Tapestation using Agilent RNA analysis screentapes (Agilent Genomics, 5067-5576). RNA with RNA Integrity Number (RIN) scores preferably > 8.0 were used in the preparation of RNA-seq libraries. At minimum, 200ng of total RNA was used in the reactions for the Illumina TruSeq stranded RNA-seq library preparation kit (Illumina, RS-122-2101) according to the manufacturer’s instructions with the modification to use Superscript III Reverse Transcriptase enzyme (Invitrogen, 18080044) during the first strand cDNA synthesis step. Completed libraries were checked for quality and average fragment size using D1000 screen tapes and molar concentration determined using NEBNext qPCR library quantification kit. Libraries were diluted to 4nM, pooled and denatured according to the instructions for Illumina NextSeq 550/500. Libraries were sequenced on the NextSeq 500 or 550 with settings for single-index, paired-end sequencing with 75 cycles per end. To analyze the data, first adapter contamination was trimmed from the reads using Trimmomatic v.0.36 (Bolger et al., 2014) and aligned to the appropriate genome assembly using the STAR aligner v.2.7.1a (Dobin et al., 2012)Alignment to hg19 used gencode.v19.annotation.gtf and alignment to mm10 used gencode.vM25.annotation.gtf. Differential gene analysis was performed in R v.3.6.3 using DESeq2 v.1.26.0 (Love et al., 2014) and surrogate variable analysis performed with the R package sva v.3.34.0 (Leek et al., 2012).

### Selection of hg19 4C viewpoint primers

Primers for 4C in HoxA region based on locations orthologous to mouse (mm9) HoxA 4C viewpoints. Primers were designed against NlaIII and DpnII cut sites that produce a fragment near or overlapping the element of interest. The primers were chosen from the 4C-seq primer database from 4cseq_pipe (https://www.wisdom.weizmann.ac.il/~atanay/4cseq_pipe/4c_primer_db_manual.pdf; (van de Werken et al., 2012) and are listed in **Supplemental Table 12**.

### 4C library preparation

Crosslinked cell pellets or crosslinked embryonic tissue was resuspended in 1x Cell lysis buffer (50mM Tris-HCl, pH 8.0, 140mM NaCl, 1mM EDTA, pH 8.0, 10% glycerol, 0.5% IGEPAL CA-630, 0.25% Triton X-100) and nuclei released in a prechilled dounce homogenizer by 10 strokes with a loose glass pestle, 20 minutes rest on ice followed by 40 strokes with a tight glass pestle (Kimble Kontes 885300-0002). The nuclei were pelleted by centrifugation at 2500xg at 4C for 5 minutes, washed twice with 1x NEBuffer 3.1 and pelleted again as before. The nuclei were resuspended in 1x NEBuffer 3.1 and permeabilized by the addition of SDS, incubated for 10 minutes at 65C with shaking at 800rpm on a ThermoMixer C (Eppendorf). The SDS was neutralized with Triton X-100 and incubated at 37C for 10 minutes with gentle rocking. Additional 10x NEBuffer 3.1 was added to adjust for the addition of SDS and Triton X-100 to restore the concentration to 1X NEBuffer 3.1 before adding 200 units of NlaIII (NEB, R0125) and incubating overnight at 37C with gentle rocking. The next morning, an additional 50 units of NlaIII was added and returned to incubate at 37C for 2 hours. Quality control samples of small volume (5ul) were taken before the first addition of NlaIII (undigested) and after the second incubation with NlaIII (digested). These quality control samples were incubated with 1ug RNAse A for 20 minutes at 37C followed by incubation with 20ug Proteinase K at 65C for 1 hour and heated to 95C for 3 minutes to reverse the crosslinks. The samples were then run on a 1% agarose/ 0.5x TAE gel to visualize the completeness of digestion. If satisfactory, the main reaction tubes were heated to 65C for 20 minutes to inactivate the restriction enzyme, take another 5ul digested control sample, and divided among three prechilled 50ml conical tubes containing ligation mix (final concentrations of each: 745ul 10x T4 ligase buffer, 745ul 10% Triton X-100, 8ul 100mg/ml BSA, 5.5 ml dH2O, 2000 units of T4 DNA ligase. The ligation reactions were incubated overnight at 15C. The next day, remove 10ul from each ligation and combine. To the digested control sample and the ligation control sample incubate with RNAse A at 37C for 20 minutes and Proteinase K at 65C for 1 hour, then heated to 95C for 3 minutes and run on a 1% agarose/0.5x TAE gel to assess successful ligation. The crosslinks in the main reactions were reversed by incubating with 300ug Proteinase K at 65C overnight. The next day, RNA and protein was removed from the reactions by incubation with 10ug RNAse A at 37C for 1 hour and 300ug proteinase K at 65C for 2 hours. The digested chromatin was extracted from the ligation reactions by adding an equal volume of premixed phenol:chloroform:isoamyl alcohol (25:24:1) (Ambion, AM9732) and centrifuging at 5000xg for 10 minutes at 4C. A second extraction of the aqueous fraction was performed in a fresh 50ml conical tube using a volume of chloroform equal to the aqueous fraction and centrifuged at 5000xg for 10 minutes at 4C. The resulting aqueous phase was transferred to a new 50ml conical tube and the chromatin precipitated by addition of 1/10 volume 3M Sodium Acetate (pH 5.2) and 2.5 volumes cold ethanol and incubating at -80C for 16-64 hours. The precipitated chromatin was pelleted by centrifugation at 10,000xg for 45min at 4C and washed once with cold 70% ethanol and centrifuged again, then air dried after removing the 70% ethanol and each dried pellet separately resuspended in 10mM Tris-HCl pH 8.0. At this point, 5ul of the ligated chromatin was saved for quality control and the concentration measured with nanodrop or Qubit (Thermo Fisher, USA). Each ligation reaction was added to the second restriction digest, containing DpnII buffer to 1x final concentration and 150 units DpnII (NEB, R0543). The digestion was incubated overnight at 37C with gentle rocking. The following day another 5ul of the digested chromatin was taken to check the quality of the second digest and run on a 1% Agarose/0.5x TAE gel alongside the ligated chromatin control. When digestion was determined to be adequate from comparing the controls, the reaction was heated to 65°C for 25 minutes to inactivate the DpnII. Each digestion reaction was transferred to a fresh 50ml conical tube with 1x T4 Ligase Buffer, water and 3ul NEB T4 ligase, mixed gently and incubated at 15°C for 4 hours. Then 1/10 volume 3M Sodium Acetate pH 5.2 and 1/1000 volume glycogen was added to facilitate DNA precipitation. Cold ethanol at 2.5 volumes was added and the reactions incubated at -80°C overnight. The next day the reactions were allowed to warm at room temperature for 30 minutes and centrifuged at 10,000xg for 45 minutes at 4°C to pellet the DNA. The supernatant was removed and the pellet washed with cold 70% ethanol and centrifuged again as before. The wash was removed and the pellet allowed to air dry. Each of the three reaction pellets were allowed to resuspend in TE for 1 hour at room temperature then pooled into one tube and purified across three Qiaquick PCR purification columns (Qiagen, 28104), eluted in the provided Elution Buffer and combined. The concentration was determined by nanodrop or Qubit. The quality of the second ligation was assessed by running ∼500ng of the purified product on a 1% agarose/0.5x TAE gel.

Amplification of specific viewpoints was done by inverse PCR with primers designed against NlaIII and DpnII cut sites that produce a fragment near or overlapping the element of interest. The amplification reaction consisted of 50ng template, Roche Long Template Buffer 1 (Roche Applied Bioscience, Sigma Aldrich, 11681834001), 200nM dNTPs (Roche Applied Bioscience, Sigma Aldrich 4829042001), 200nM each of forward and reverse primer for the specific viewpoint and 0.7ul Long Template Polymerase Mix. To reduce amplification bias, eight to 16 replicate PCR reactions were performed per viewpoint. The replicate reactions were pooled and used in the subsequent indexing PCR reaction. Pooled replicate viewpoint amplification reactions were purified using QIAquick columns according to the manufacturer’s instructions, quantified by nanodrop or Qubit and five to eight indexing PCR reactions were set up per viewpoint. The indexing PCR reaction used 50-100ng template, Roche Long Template Buffer 1, 200nM dNTPs, 200nM i5 and i7 index primers and 0.7ul Long Template Polymerase Mix. The indexing primers carrying sequences recognized as [truseq 701-…, 501-…] are listed in Supplemental Table. The replicate indexing reactions were pooled and QiaQuick columns followed by Axygen bead cleanup with 1:1 ratio AxyPrep Mag PCR Clean Up beads (Axygen, 14223151). The indexed libraries, now in their final form, were checked for quality using the Agilent Tapestation (Agilent, USA) with D1000 Screen Tape. The molar library concentration was assessed using NEBNext Library Quant Kit for Illumina (NEB, E7630). Libraries were diluted to 4nM, pooled and denatured for sequencing on the Illumina NextSeq 500 or 550 as directed and sequenced using the settings for 75bp single-end, dual index sequencing. Data was processed as described at https://github.com/cotneylab/Mouse-HOXA-4C-Seq. Briefly, the biological samples were first demultiplexed by viewpoint using cutadapt (Martin, 2011), aligned to the genome using bowtie2 and analyzed using a version of r3Cseq (Thongjuea et al., 2013)) modified to allow visualization of plots over larger distances (modified version available at https://github.com/cotneylab/r3Cseq).

### Animal husbandry, tissue collection and imaging of palates

Male and female heterozygous for deletion of chr6:50,673,614-51,196,805 (ΔGCR/+) on c57bl/6 background were provided by Cyagen Animal Model Services (Santa Clara, CA, USA).

Founders were backcrossed to WT c57bl/6J mice ordered directly from Jackson Laboratories (Bar Harbor, ME, USA) and four generations of backcrossing ΔGCR/+ males to WT females was performed to reduce phenotypes due to off-target CRISPR effects. Pharyngeal arches from E11.5 embryos were collected as follows: timed matings were set up and the mice separated the next day and females checked for plugs. Noon of that day was counted as day 0.5. On day 11.5, mice with noted plugs were euthanized with CO2 in accordance with the AVMA Guidelines for the Euthanasia of Animals. Uterine horns were removed from the euthanized female and rinsed in PBS, individual embryos were removed from the uterine sacs and individually dissected for pharyngeal arches, limb buds, heart and forebrain. The dissected tissue was frozen on a dry ice/ethanol slurry and stored at -80°C until use. The remainder of the body was used for genotyping. DNA for genotyping was extracted using a high salt lysis buffer with proteinase K followed by chloroform extraction. Primers spanning the deletion were used to confirm genotype (sequences in **Supplemental Table 11**). For initial assessment of palatal morphology and fetal weight, timed matings were set up between ΔGCR/+ mice. The mice were separated the following morning and noon of that day was counted as day 0.5. At day 17.5, the female mice were checked for pregnancy and euthanized with CO2 in accordance with the AVMA Guidelines for the Euthanasia of Animals. The uterine horns were removed from the euthanized mother, rinsed in cold PBS, the individual pups removed from the uterine sacs, rinsed in PBS, gently dried and weighed, a tail clip taken for genotyping and the body fixed in neutral buffered formalin for at least 48h prior to removal of upper jaw for palate photographs. For soft tissue and skeletal imaging, timed matings were similarly set up and embryos taken at E18.5 as described above. Embryos were fixed in neutral buffered formalin for 24 hours and transferred to 70% ethanol prior to shipping for analysis. All protocols for animal use and care were approved by the University of Connecticut Institutional Animal Care and Use Committee and conform to the NIH Guide for the Care and Use of Laboratory Animals. CO2 euthanasia was performed in accordance with the AVMA Guidelines for the Euthanasia of Animals.

### Microcomputed tomography imaging and analysis of embryos

In preparation for microcomputed tomography (microCT) imaging, fixed E18.5 embryos were briefly rinsed in 70% ethanol and placed individually on custom styrofoam beds that held the embryos in an upright position. Embryos were then individually scanned using a Skyscan model 1275 benchtop microCT (Bruker BioSpin Corporation) using the following parameters: an isotropic resolution of 18 microns, 40 kV, 180 microAmp, 45 ms exposure, 0.3° rotation step, 180° rotation, and 4 frame averaging. No filter was used. 16-bit raw images from all scans were reconstructed to 8-bit bmp files using NRecon software v1.7.4.2 (Bruker BioSpin Corporation). Reconstructed scan data were then imported into Drishti volume exploration software (version 2.63) (Limaye, 2012) for 3D rendering. Separate rendering settings were optimized for visualization and phenotypic assessment of both mineralized and soft tissues.

To visualize the pterygoid/basisphenoid and tympanic regions in isolation, the majority of the craniofacial bone around these structures was masked using clipping planes. Then, the morphological operations (MOP) carve function was used to remove remaining bone from around the complex. To make rotational movies of the complexes to aid their inspection, the Keyframe Editor function of Drishti was employed. For this, a new rotational axis was assigned for each volume and the initial keyframe set to mark the starting view of the rotation. The desired end of the rotation was set using the Bricks Editor function and a second keyframe set. All interpolated keyframes between the starting and ending keyframe were then saved as an image sequence in png format. Image sequences were then opened in Adobe Photoshop 2021 and rendered in mp4 format. Selected images from the renderings were saved and optimized for contrast, color, and background using Adobe Photoshop.

### Copy number detection through shallow whole-genome sequencing

Fetal gDNA was extracted and isolated upon biopsy from fetal tissue on QIAcube, using the QIAamp DNA Blood Mini Kit (Qiagen). Shallow whole genome sequencing for copy number variant analysis was performed on extracted gDNA using the NEXTflex Rapid DNA Sequencing kit (Bioo Scientific). The normalized libraries were sequenced on a HiSeq 3000 platform (Illumina) and data-analysis was performed according to Raman *et al*. (Raman et al., 2018) with a 400 kb resolution.

### Mendeliome analysis

Extracted fetal gDNA was further used to perform whole exome sequencing (WES). The coding exons and flanking intronic regions were enriched with the SureSelectXT Low Input All Exon v7 kit (Agilent Technologies) followed by dual index, paired-end (2 × 150 bp) sequencing on a NovaSeq 6000 platform (Illumina). Raw sequence reads were processed using an in-house developed pipeline. Reads were aligned to GRCh38/hg38 and data analysis was limited to the at that moment applicable Mendeliome panel (version 2), containing 4007 genes related to a known disease. At least 90% of the coding regions of the included genes had a minimum coverage of 20x. A trio-analysis was performed. Only variants classified as class 4 (possibly pathogenic) and class 5 (pathogenic), if clinically relevant, were reported.

### Use of publicly available datasets

The following datasets were downloaded from the Gene Expression Omnibus (GEO): Series GSE105028: “Architectural proteins and pluripotency factors cooperate to orchestrate the transcriptional response of hESCs to temperature stress” (Lyu et al., 2018); Series GSE145327: “CTCF ChIP-seq in undifferentiated H9 hESCs and H9-derived CNCCs” (Long et al., 2020) and Series GSE104173: “Expression data from retinoic acid-induced differentiation of human embryonic stem cells (hESCs)”. Plots generated from HiC data using the HiC Browser hosted at Northwestern University (3dgenome.fsm.northwestern.edu/view.php) included Adrenal Gland, Cortex, Right Ventricle (Schmitt et al., 2016); GM12878, IMR90 (Rao et al., 2014) and Liver (Leung et al., 2015).

### Code availability

Code used to analyze 4C-seq data is available at github.com/cotney/Mouse-HOXA-4C-Seq. The most recent HiC pipeline is at github.com/awilderman/HiC. Code used in the analysis of human and mouse superenhancers is available at github.com/awilderman/Thesis.

## Notes

https://cotney.research.uchc.edu/hoxa_gcr

### Competing Interest Statement

The authors have declared no competing interest.

### Summary of Updates

Corrected author list

## References

Ahituv, N., Zhu, Y., Visel, A., Holt, A., Afzal, V., Pennacchio, L.A., and Rubin, E.M. (2007). Deletion of ultraconserved elements yields viable mice. PLoS Biol. 5, e234.

Alasti, F., Sadeghi, A., Sanati, M.H., Farhadi, M., Stollar, E., Somers, T., and Van Camp, G. (2008). A mutation in HOXA2 is responsible for autosomal-recessive microtia in an Iranian family. Am. J. Hum. Genet. 82, 982–991.

Attanasio, C., Nord, A.S., Zhu, Y., Blow, M.J., Li, Z., Liberton, D.K., Morrison, H., Plajzer-Frick, I., Holt, A., Hosseini, R., et al. (2013). Fine tuning of craniofacial morphology by distant-acting enhancers. Science 342, 1241006.

Aydin, E., Kloos, D.-P., Gay, E., Jonker, W., Hu, L., Bullwinkel, J., Brown, J.P., Manukyan, M., Giera, M., Singh, P.B., et al. (2015). A hypomorphic Cbx3 allele causes prenatal growth restriction and perinatal energy homeostasis defects. J. Biosci. 40, 325–338.

Barrow, J.R., and Capecchi, M.R. (1999). Compensatory defects associated with mutations in Hoxa1 restore normal palatogenesis to Hoxa2 mutants. Development 126, 5011–5026.

Belaghzal, H., Dekker, J., and Gibcus, J.H. (2017). Hi-C 2.0: An optimized Hi-C procedure for high-resolution genome-wide mapping of chromosome conformation. Methods 123, 56–65.

Berlivet, S., Paquette, D., Dumouchel, A., Langlais, D., Dostie, J., and Kmita, M. (2013). Clustering of tissue-specific sub-TADs accompanies the regulation of HoxA genes in developing limbs. PLoS Genet. 9, e1004018.

Blobel, G.A., Higgs, D.R., Mitchell, J.A., Notani, D., and Young, R.A. (2021). Testing the super-enhancer concept. Nat. Rev. Genet. 22, 749–755.

Bolger, A.M., Lohse, M., and Usadel, B. (2014). Trimmomatic: a flexible trimmer for Illumina sequence data. Bioinformatics 30, 2114–2120.

Bolt, C.C., and Duboule, D. (2020). The regulatory landscapes of developmental genes. Development 147.

Bonfante, B., Faux, P., Navarro, N., Mendoza-Revilla, J., Dubied, M., Montillot, C., Wentworth, E., Poloni, L., Varón-González, C., Jones, P., et al. (2021). A GWAS in Latin Americans identifies novel face shape loci, implicating VPS13B and a Denisovan introgressed region in facial variation. Sci Adv 7.

Choi, J., Lysakovskaia, K., Stik, G., Demel, C., Söding, J., Tian, T.V., Graf, T., and Cramer, P. (2021). Evidence for additive and synergistic action of mammalian enhancers during cell fate determination. Elife 10.

Dave, K., Sur, I., Yan, J., Zhang, J., Kaasinen, E., Zhong, F., Blaas, L., Li, X., Kharazi, S., Gustafsson, C., et al. (2017). Mice deficient of Myc super-enhancer region reveal differential control mechanism between normal and pathological growth. Elife 6, e23382.

Dickel, D.E., Ypsilanti, A.R., Pla, R., Zhu, Y., Barozzi, I., Mannion, B.J., Khin, Y.S., Fukuda-Yuzawa, Y., Plajzer-Frick, I., Pickle, C.S., et al. (2018). Ultraconserved Enhancers Are Required for Normal Development. Cell 172, 491–499.e15.

Dixon, J.R., Selvaraj, S., Yue, F., Kim, A., Li, Y., Shen, Y., Hu, M., Liu, J.S., and Ren, B. (2012). Topological domains in mammalian genomes identified by analysis of chromatin interactions. Nature 485, 376–380.

Dobin, A., Davis, C.A., Schlesinger, F., Drenkow, J., Zaleski, C., Jha, S., Batut, P., Chaisson, M., and Gingeras, T.R. (2012). STAR: ultrafast universal RNA-seq aligner. Bioinformatics 29, 15–21.

Donaldson, I.J., Amin, S., Hensman, J.J., Kutejova, E., Rattray, M., Lawrence, N., Hayes, A., Ward, C.M., and Bobola, N. (2012). Genome-wide occupancy links Hoxa2 to Wnt-β-catenin signaling in mouse embryonic development. Nucleic Acids Res. 40, 3990–4001.

Etchevers, H.C., Amiel, J., and Lyonnet, S. (2006). Molecular bases of human neurocristopathies. Adv. Exp. Med. Biol. 589, 213–234.

Farooq, U., Saravanan, B., Islam, Z., Walavalkar, K., Singh, A.K., Jayani, R.S., Meel, S., Swaminathan, S., and Notani, D. (2021). An interdependent network of functional enhancers regulates transcription and EZH2 loading at the INK4a/ARF locus. Cell Rep. 34, 108898.

Firth, H.V., Richards, S.M., Bevan, A.P., Clayton, S., Corpas, M., Rajan, D., Van Vooren, S., Moreau, Y., Pettett, R.M., and Carter, N.P. (2009). DECIPHER: Database of Chromosomal Imbalance and Phenotype in Humans Using Ensembl Resources. Am. J. Hum. Genet. 84, 524–533.

Franke, M., Ibrahim, D.M., Andrey, G., Schwarzer, W., Heinrich, V., Schöpflin, R., Kraft, K., Kempfer, R., Jerković, I., Chan, W.-L., et al. (2016). Formation of new chromatin domains determines pathogenicity of genomic duplications. Nature 538, 265–269.

Fulco, C.P., Munschauer, M., Anyoha, R., Munson, G., Grossman, S.R., Perez, E.M., Kane, M., Cleary, B., Lander, E.S., and Engreitz, J.M. (2016). Systematic mapping of functional enhancer-promoter connections with CRISPR interference. Science 354, 769–773.

Gendron-Maguire, M., Mallo, M., Zhang, M., and Gridley, T. (1993). Hoxa-2 mutant mice exhibit homeotic transformation of skeletal elements derived from cranial neural crest. Cell 75, 1317–1331.

Gillentine, M.A., Wang, T., Hoekzema, K., Rosenfeld, J., Liu, P., Guo, H., Kim, C.N., De Vries, B.B.A., Vissers, L.E.L.M., Nordenskjold, M., et al. (2021). Rare deleterious mutations of HNRNP genes result in shared neurodevelopmental disorders. Genome Med. 13, 63.

Guerrero, L., Marco-Ferreres, R., Serrano, A.L., Arredondo, J.J., and Cervera, M. (2010). Secondary enhancers synergise with primary enhancers to guarantee fine-tuned muscle gene expression. Dev. Biol. 337, 16–28.

Guo, Y., Xu, Q., Canzio, D., Shou, J., Li, J., Gorkin, D.U., Jung, I., Wu, H., Zhai, Y., Tang, Y., et al. (2015). CRISPR Inversion of CTCF Sites Alters Genome Topology and Enhancer/Promoter Function. Cell 162, 900–910.

Hay, D., Hughes, J.R., Babbs, C., Davies, J.O.J., Graham, B.J., Hanssen, L.L.P., Kassouf, M.T., Oudelaar, A.M., Sharpe, J.A., Suciu, M.C., et al. (2016). Genetic dissection of the α-globin super-enhancer in vivo. Nat. Genet. 48, 895–903.

Hnisz, D., Abraham, B.J., Lee, T.I., Lau, A., Saint-André, V., Sigova, A.A., Hoke, H.A., and Young, R.A. (2013). Super-enhancers in the control of cell identity and disease. Cell 155, 934–947.

Hnisz, D., Weintraub, A.S., Day, D.S., Valton, A.-L., Bak, R.O., Li, C.H., Goldmann, J., Lajoie, B.R., Fan, Z.P., Sigova, A.A., et al. (2016). Activation of proto-oncogenes by disruption of chromosome neighborhoods. Science 351, 1454–1458.

Huang, J., Li, K., Cai, W., Liu, X., Zhang, Y., Orkin, S.H., Xu, J., and Yuan, G.-C. (2018). Dissecting super-enhancer hierarchy based on chromatin interactions. Nat. Commun. 9, 943.

Hunt, P., Gulisano, M., Cook, M., Sham, M.H., Faiella, A., Wilkinson, D., Boncinelli, E., and Krumlauf, R. (1991). A distinct Hox code for the branchial region of the vertebrate head. Nature 353, 861–864.

Khan, A., and Zhang, X. (2016). dbSUPER: a database of super-enhancers in mouse and human genome. Nucleic Acids Res. 44, D164–D171.

Khoury, A., Achinger-Kawecka, J., Bert, S.A., Smith, G.C., French, H.J., Luu, P.-L., Peters, T.J., Du, Q., Parry, A.J., Valdes-Mora, F., et al. (2020). Constitutively bound CTCF sites maintain 3D chromatin architecture and long-range epigenetically regulated domains. Nat. Commun. 11, 54.

Kim, H.J., Kim, N.C., Wang, Y.-D., Scarborough, E.A., Moore, J., Diaz, Z., MacLea, K.S., Freibaum, B., Li, S., Molliex, A., et al. (2013). Mutations in prion-like domains in hnRNPA2B1 and hnRNPA1 cause multisystem proteinopathy and ALS. Nature 495, 467–473.

Kleinjan, D.A., and Lettice, L.A. (2008). Long-range gene control and genetic disease. Adv. Genet. 61, 339–388.

Kmita, M., Tarchini, B., Zàkàny, J., Logan, M., Tabin, C.J., and Duboule, D. (2005). Early developmental arrest of mammalian limbs lacking HoxA/HoxD gene function. Nature 435, 1113–1116.

Lee, A.P., Koh, E.G.L., Tay, A., Brenner, S., and Venkatesh, B. (2006). Highly conserved syntenic blocks at the vertebrate Hox loci and conserved regulatory elements within and outside Hox gene clusters. Proc. Natl. Acad. Sci. U. S. A. 103, 6994–6999.

Leek, J.T., Johnson, W.E., Parker, H.S., Jaffe, A.E., and Storey, J.D. (2012). The sva package for removing batch effects and other unwanted variation in high-throughput experiments. Bioinformatics 28, 882–883.

Lettice, L.A., Heaney, S.J.H., Purdie, L.A., Li, L., de Beer, P., Oostra, B.A., Goode, D., Elgar, G., Hill, R.E., and de Graaff, E. (2003). A long-range Shh enhancer regulates expression in the developing limb and fin and is associated with preaxial polydactyly. Hum. Mol. Genet. 12, 1725–1735.

Leung, A.W., Murdoch, B., Salem, A.F., Prasad, M.S., Gomez, G.A., and García-Castro, M.I. (2016). WNT/β-catenin signaling mediates human neural crest induction via a pre-neural border intermediate. Development 143, 398–410.

Leung, D., Jung, I., Rajagopal, N., Schmitt, A., Selvaraj, S., Lee, A.Y., Yen, C.-A., Lin, S., Lin, Y., Qiu, Y., et al. (2015). Integrative analysis of haplotype-resolved epigenomes across human tissues. Nature 518, 350–354.

Liao, Y., Wang, J., Jaehnig, E.J., Shi, Z., and Zhang, B. (2019). WebGestalt 2019: gene set analysis toolkit with revamped UIs and APIs. Nucleic Acids Res. 47, W199–W205.

Limaye, A. (2012). Drishti: a volume exploration and presentation tool. In Proceedings Volume 8506, Developments in X-Ray Tomography VIII; 85060X (2012),.

Long, H.K., Osterwalder, M., Welsh, I.C., Hansen, K., Davies, J.O.J., Liu, Y.E., Koska, M., Adams, A.T., Aho, R., Arora, N., et al. (2020). Loss of Extreme Long-Range Enhancers in Human Neural Crest Drives a Craniofacial Disorder. Cell Stem Cell 27, 765–783.e14.

Love, M.I., Huber, W., and Anders, S. (2014). Moderated estimation of fold change and dispersion for RNA-seq data with DESeq2. Genome Biol. 15, 550.

Lovén, J., Hoke, H.A., Lin, C.Y., Lau, A., Orlando, D.A., Vakoc, C.R., Bradner, J.E., Lee, T.I., and Young, R.A. (2013). Selective Inhibition of Tumor Oncogenes by Disruption of Super-Enhancers. Cell 153, 320–334.

Lupiáñez, D.G., Kraft, K., Heinrich, V., Krawitz, P., Brancati, F., Klopocki, E., Horn, D., Kayserili, H., Opitz, J.M., Laxova, R., et al. (2015). Disruptions of topological chromatin domains cause pathogenic rewiring of gene-enhancer interactions. Cell 161, 1012–1025.

Lyu, X., Rowley, M.J., and Corces, V.G. (2018). Architectural Proteins and Pluripotency Factors Cooperate to Orchestrate the Transcriptional Response of hESCs to Temperature Stress. Mol. Cell 71, 940–955.e7.

Martin, M. (2011). Cutadapt removes adapter sequences from high-throughput sequencing reads. EMBnet.journal 17, 10–12.

Melo, U.S., Schöpflin, R., Acuna-Hidalgo, R., Mensah, M.A., Fischer-Zirnsak, B., Holtgrewe, M., Klever, M.-K., Türkmen, S., Heinrich, V., Pluym, I.D., et al. (2020). Hi-C Identifies Complex Genomic Rearrangements and TAD-Shuffling in Developmental Diseases. Am. J. Hum. Genet. 106, 872–884.

Melo, U.S., Piard, J., Fischer-Zirnsak, B., Klever, M.-K., Schöpflin, R., Mensah, M.A., Holtgrewe, M., Arbez-Gindre, F., Martin, A., Guigue, V., et al. (2021). Complete lung agenesis caused by complex genomic rearrangements with neo-TAD formation at the SHH locus. Hum. Genet. 140, 1459–1469.

Middelkamp, S., Vlaar, J.M., Giltay, J., Korzelius, J., Besselink, N., Boymans, S., Janssen, R., de la Fonteijne, L., van Binsbergen, E., van Roosmalen, M.J., et al. (2019). Prioritization of genes driving congenital phenotypes of patients with de novo genomic structural variants.

Minoux, M., and Rijli, F.M. (2010). Molecular mechanisms of cranial neural crest cell migration and patterning in craniofacial development. Development 137, 2605–2621.

Moorthy, S.D., Davidson, S., Shchuka, V.M., Singh, G., Malek-Gilani, N., Langroudi, L., Martchenko, A., So, V., Macpherson, N.N., and Mitchell, J.A. (2017). Enhancers and super-enhancers have an equivalent regulatory role in embryonic stem cells through regulation of single or multiple genes. Genome Res. 27, 246–258.

Nanni, L., Ceri, S., and Logie, C. (2020). Spatial patterns of CTCF sites define the anatomy of TADs and their boundaries. Genome Biol. 21, 197.

Nóbrega, M.A., Zhu, Y., Plajzer-Frick, I., Afzal, V., and Rubin, E.M. (2004). Megabase deletions of gene deserts result in viable mice. Nature 431, 988–993.

Nolte, M.J., Wang, Y., Deng, J.M., Swinton, P.G., Wei, C., Guindani, M., Schwartz, R.J., and Behringer, R.R. (2014). Functional analysis of limb transcriptional enhancers in the mouse. Evol. Dev. 16, 207–223.

Osterwalder, M., Barozzi, I., Tissières, V., Fukuda-Yuzawa, Y., Mannion, B.J., Afzal, S.Y., Lee, E.A., Zhu, Y., Plajzer-Frick, I., Pickle, C.S., et al. (2018). Enhancer redundancy provides phenotypic robustness in mammalian development. Nature 554, 239–243.

Pott, S., and Lieb, J.D. (2015). What are super-enhancers? Nat. Genet. 47, 8–12.

Prabhakar, S., Visel, A., Akiyama, J.A., Shoukry, M., Lewis, K.D., Holt, A., Plajzer-Frick, I., Morrison, H., Fitz Patrick, D.R., Afzal, V., et al. (2008). Human-Specific Gain of Function in a Developmental Enhancer. Science 321.

Qi, X., Pang, Q., Wang, J., Zhao, Z., Wang, O., Xu, L., Mao, J., Jiang, Y., Li, M., Xing, X., et al. (2017). Familial Early-Onset Paget’s Disease of Bone Associated with a Novel hnRNPA2B1 Mutation. Calcif. Tissue Int. 101, 159–169.

Quinlan, A.R., and Hall, I.M. (2010). BEDTools: a flexible suite of utilities for comparing genomic features. Bioinformatics 26, 841–842.

Quinonez, S.C., and Innis, J.W. (2014). Human HOX gene disorders. Mol. Genet. Metab. 111, 4–15.

Raman, L., Dheedene, A., De Smet, M., Van Dorpe, J., and Menten, B. (2018). WisecondorX: improved copy number detection for routine shallow whole-genome sequencing. Nucleic Acids Res. 47, 1605–1614.

Ramírez, F., Bhardwaj, V., Arrigoni, L., Lam, K.C., Grüning, B.A., Villaveces, J., Habermann, B., Akhtar, A., and Manke, T. (2018). High-resolution TADs reveal DNA sequences underlying genome organization in flies. Nat. Commun. 9, 189.

Rao, S.S.P., Huntley, M.H., Durand, N.C., Stamenova, E.K., Bochkov, I.D., Robinson, J.T., Sanborn, A.L., Machol, I., Omer, A.D., Lander, E.S., et al. (2014). A 3D map of the human genome at kilobase resolution reveals principles of chromatin looping. Cell 159, 1665–1680.

Rijli, F.M., Mark, M., Lakkaraju, S., Dierich, A., Doll, P., de G, P.C.L., Mol, I., and de Biologie, M. (1993). A Homeotic Transformation Is Generated in the Rostra1 Branchial Region of the Head by Disruption of Hoxa-2, Which Acts as a Selector Gene. Cell 75, 1333–1349.

Santagati, F., Minoux, M., Ren, S.-Y., and Rijli, F.M. (2005). Temporal requirement of Hoxa2 in cranial neural crest skeletal morphogenesis. Development 132, 4927–4936.

Saravanan, B., Soota, D., Islam, Z., Majumdar, S., Mann, R., Meel, S., Farooq, U., Walavalkar, K., Gayen, S., Singh, A.K., et al. (2020). Ligand dependent gene regulation by transient ERα clustered enhancers. PLoS Genet. 16, e1008516.

Schmitt, A.D., Hu, M., Jung, I., Xu, Z., Qiu, Y., Tan, C.L., Li, Y., Lin, S., Lin, Y., Barr, C.L., et al. (2016). A Compendium of Chromatin Contact Maps Reveals Spatially Active Regions in the Human Genome. Cell Rep. 17, 2042–2059.

Servant, N., Varoquaux, N., Lajoie, B.R., Viara, E., Chen, C.-J., Vert, J.-P., Heard, E., Dekker, J., and Barillot, E. (2015). HiC-Pro: an optimized and flexible pipeline for Hi-C data processing. Genome Biol. 16, 259.

Spitz, F., and Furlong, E.E.M. (2012). Transcription factors: from enhancer binding to developmental control. Nat. Rev. Genet. 13, 613–626.

Spitz, F., Gonzalez, F., and Duboule, D. (2003). A global control region defines a chromosomal regulatory landscape containing the HoxD cluster. Cell 113, 405–417.

Symmons, O., Pan, L., Remeseiro, S., Aktas, T., Klein, F., Huber, W., and Spitz, F. (2016). The Shh Topological Domain Facilitates the Action of Remote Enhancers by Reducing the Effects of Genomic Distances. Dev. Cell 39, 529–543.

Thongjuea, S., Stadhouders, R., Grosveld, F.G., Soler, E., and Lenhard, B. (2013). r3Cseq: an R/Bioconductor package for the discovery of long-range genomic interactions from chromosome conformation capture and next-generation sequencing data. Nucleic Acids Res. 41, e132.

Tischfield, M.A., Bosley, T.M., Salih, M.A.M., Alorainy, I.A., Sener, E.C., Nester, M.J., Oystreck, D.T., Chan, W.-M., Andrews, C., Erickson, R.P., et al. (2005). Homozygous HOXA1 mutations disrupt human brainstem, inner ear, cardiovascular and cognitive development. Nat. Genet. 37, 1035–1037.

VanOudenhove, J., Yankee, T.N., Wilderman, A., and Cotney, J. (2020). Epigenomic and Transcriptomic Dynamics During Human Heart Organogenesis. Circ. Res. 127, e184–e209.

Vietri Rudan, M., Barrington, C., Henderson, S., Ernst, C., Odom, D.T., Tanay, A., and Hadjur, S. (2015). Comparative Hi-C reveals that CTCF underlies evolution of chromosomal domain architecture. Cell Rep. 10, 1297–1309.

Visel, A., Minovitsky, S., Dubchak, I., and Pennacchio, L.A. (2007). VISTA Enhancer Browser--a database of tissue-specific human enhancers. Nucleic Acids Res. 35, D88–D92.

Visel, A., Prabhakar, S., Akiyama, J.A., Shoukry, M., Lewis, K.D., Holt, A., Plajzer-Frick, I., Afzal, V., Rubin, E.M., and Pennacchio, L.A. (2008). Ultraconservation identifies a small subset of extremely constrained developmental enhancers. Nat. Genet. 40, 158–160.

Weischenfeldt, J., Dubash, T., Drainas, A.P., Mardin, B.R., Chen, Y., Stütz, A.M., Waszak, S.M., Bosco, G., Halvorsen, A.R., Raeder, B., et al. (2017). Pan-cancer analysis of somatic copy-number alterations implicates IRS4 and IGF2 in enhancer hijacking. Nat. Genet. 49, 65–74.

van de Werken, H.J.G., Landan, G., Holwerda, S.J.B., Hoichman, M., Klous, P., Chachik, R., Splinter, E., Valdes-Quezada, C., Oz, Y., Bouwman, B.A.M., et al. (2012). Robust 4C-seq data analysis to screen for regulatory DNA interactions. Nat. Methods 9, 969–972.

White, J.D., Indencleef, K., Naqvi, S., Eller, R.J., Hoskens, H., Roosenboom, J., Lee, M.K., Li, J., Mohammed, J., Richmond, S., et al. (2021). Insights into the genetic architecture of the human face. Nat. Genet. 53, 45–53.

Whyte, W.A., Orlando, D.A., Hnisz, D., Abraham, B.J., Lin, C.Y., Kagey, M.H., Rahl, P.B., Lee, T.I., and Young, R.A. (2013). Master Transcription Factors and Mediator Establish Super-Enhancers at Key Cell Identity Genes. Cell 153, 307–319.

Wilderman, A., VanOudenhove, J., Kron, J., Noonan, J.P., and Cotney, J. (2018). High-Resolution Epigenomic Atlas of Human Embryonic Craniofacial Development. Cell Rep. 23, 1581–1597.

Woltering, J.M., Noordermeer, D., Leleu, M., and Duboule, D. (2014). Conservation and divergence of regulatory strategies at Hox Loci and the origin of tetrapod digits. PLoS Biol. 12, e1001773.

Yankee, T.N., Wilderman, A., Winchester, E.W., VanOudenhove, J., and Cotney, J. (2022). Integrative analysis of transcriptomics in human craniofacial development reveals novel candidate disease genes.

Yu, G., Wang, L.G., Han, Y., and He, Q.Y. (2012). clusterProfiler: an R package for comparing biological themes among gene clusters. OMICS.

Zhang, B., Kirov, S., and Snoddy, J. (2005). WebGestalt: an integrated system for exploring gene sets in various biological contexts. Nucleic Acids Res. 33, W741–W748.

Zhang, X., Jeong, M., Huang, X., Wang, X.Q., Wang, X., Zhou, W., Shamim, M.S., Gore, H., Himadewi, P., Liu, Y., et al. (2020). Large DNA Methylation Nadirs Anchor Chromatin Loops Maintaining Hematopoietic Stem Cell Identity. Mol. Cell 78, 506–521.e6.

