## Supplemental Text and Figures for "A distant global control region is essential for normal expression of anterior *HOXA* genes during mouse and human craniofacial development"

*Gynecologists report:*
A 30-year old G1P0 was referred to our tertiary centre for targeted ultrasonography at 13 weeks due to abnormal profile. We noted an abnormal retronasal triangle, absence of the palate in sagittal view and a protruding median part of the maxilla (‘parrot beak’), indicating a severe bilateral cheilognathopalatoschisis. Forearms and hands were structurally normal but afunctional. Both feet were abnormally positioned. Termination of pregnancy on the couple's request was performed at 14 weeks. Consent was given to publish photographs and molecular data.

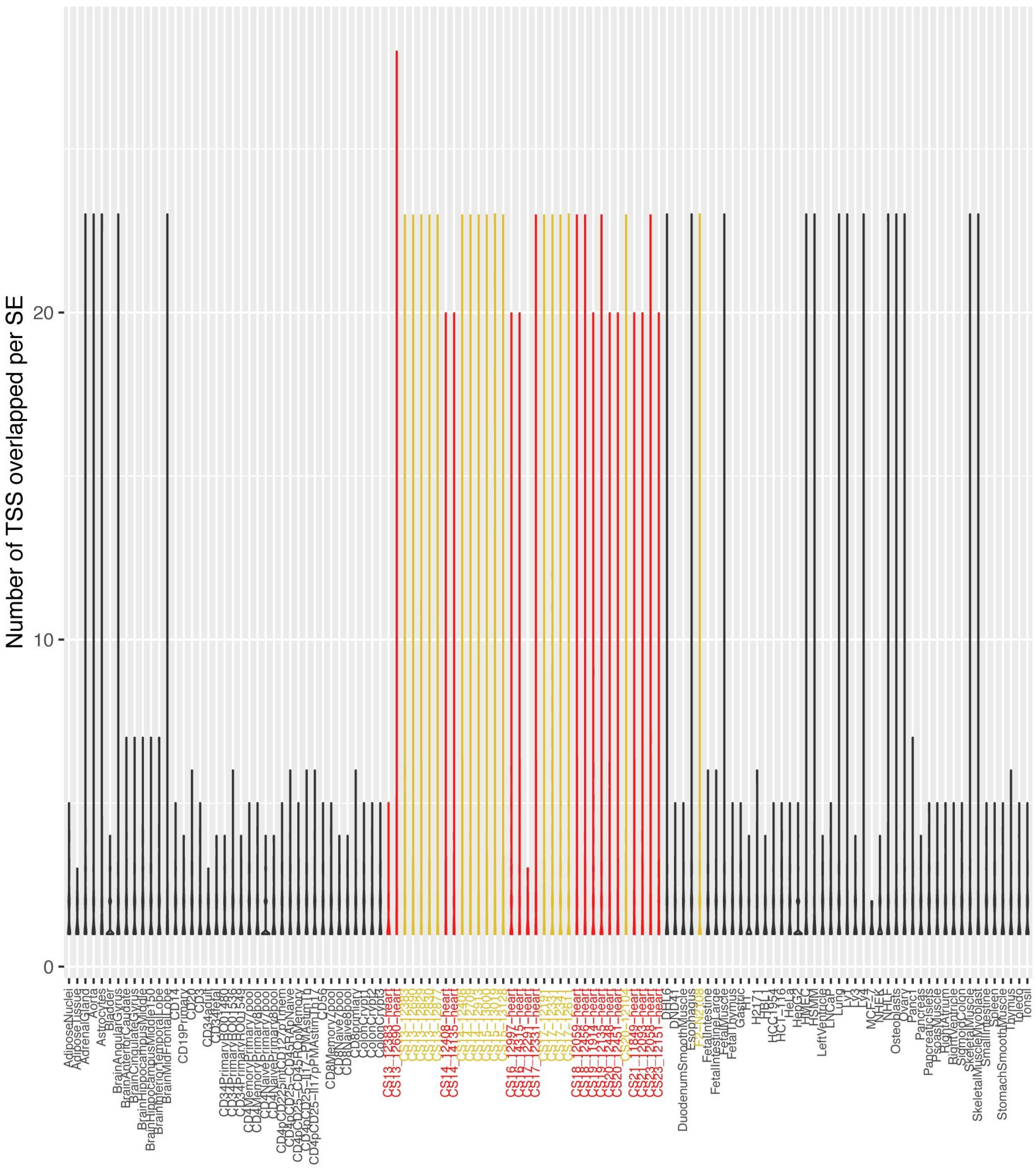


**Figure 1 Supplement 1-** Violin plots of number of transcription start sites (TSSs) overlapped by superenhancers in various tissues. Plots in black reference tissues and cell types in the dbSuper database. Plots in yellow are the results from human embryonic craniofacial tissue. Plots in red are the results from human embryonic heart. Superenhancers encompassing at minimum 1 and as many as 23 TSSs are found across multiple tissues. Median number of TSSs for protein-coding genes encompassed by superenhancers in dbSuper tissues and cell lines = 1. Median number of TSSs for protein coding genes encompassed by superenhancers in craniofacial and heart samples ranged from 1-3.

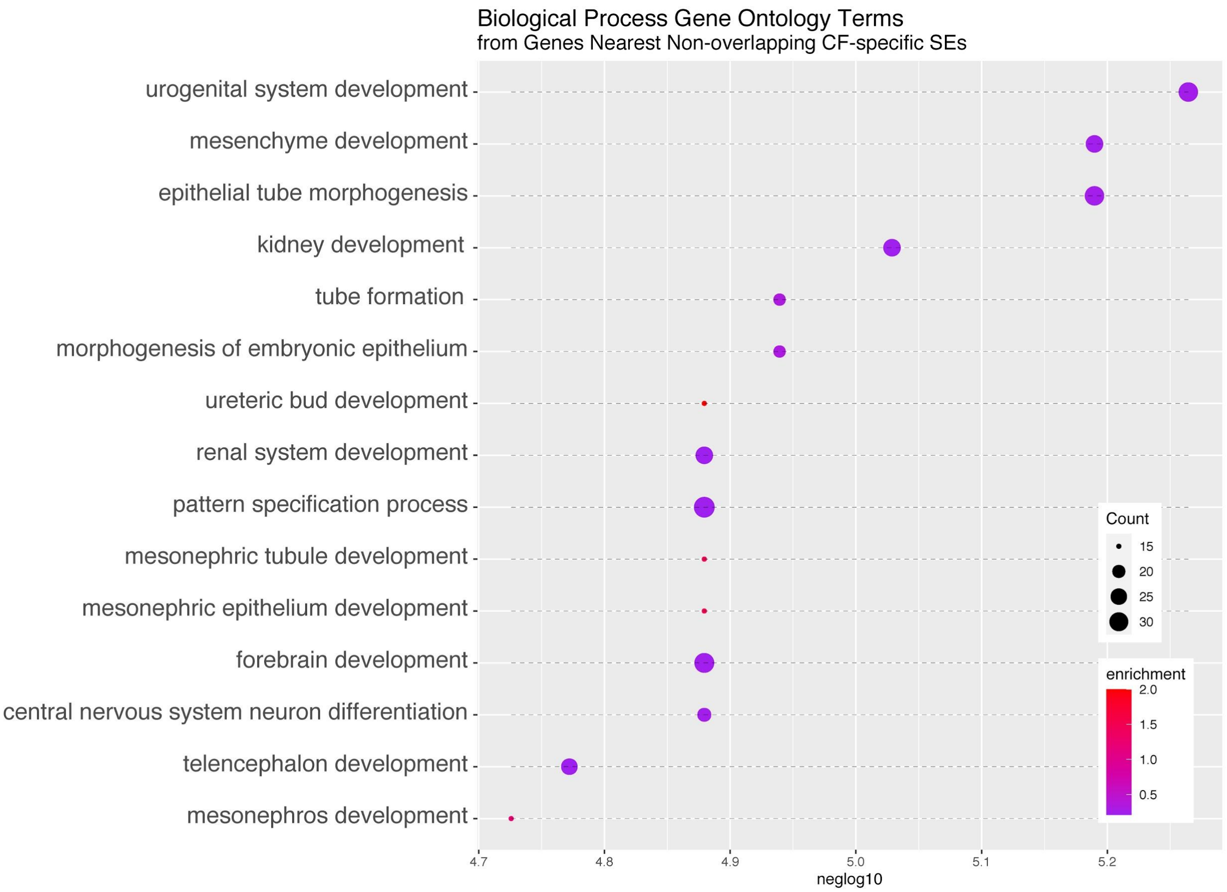


**Figure 1 Supplement 2-** Biological Process Gene Ontology terms enriched in genes assigned to superenhancers in non-coding regions. Assignment of the two nearest genes was done through the BedTools suite function "closest".

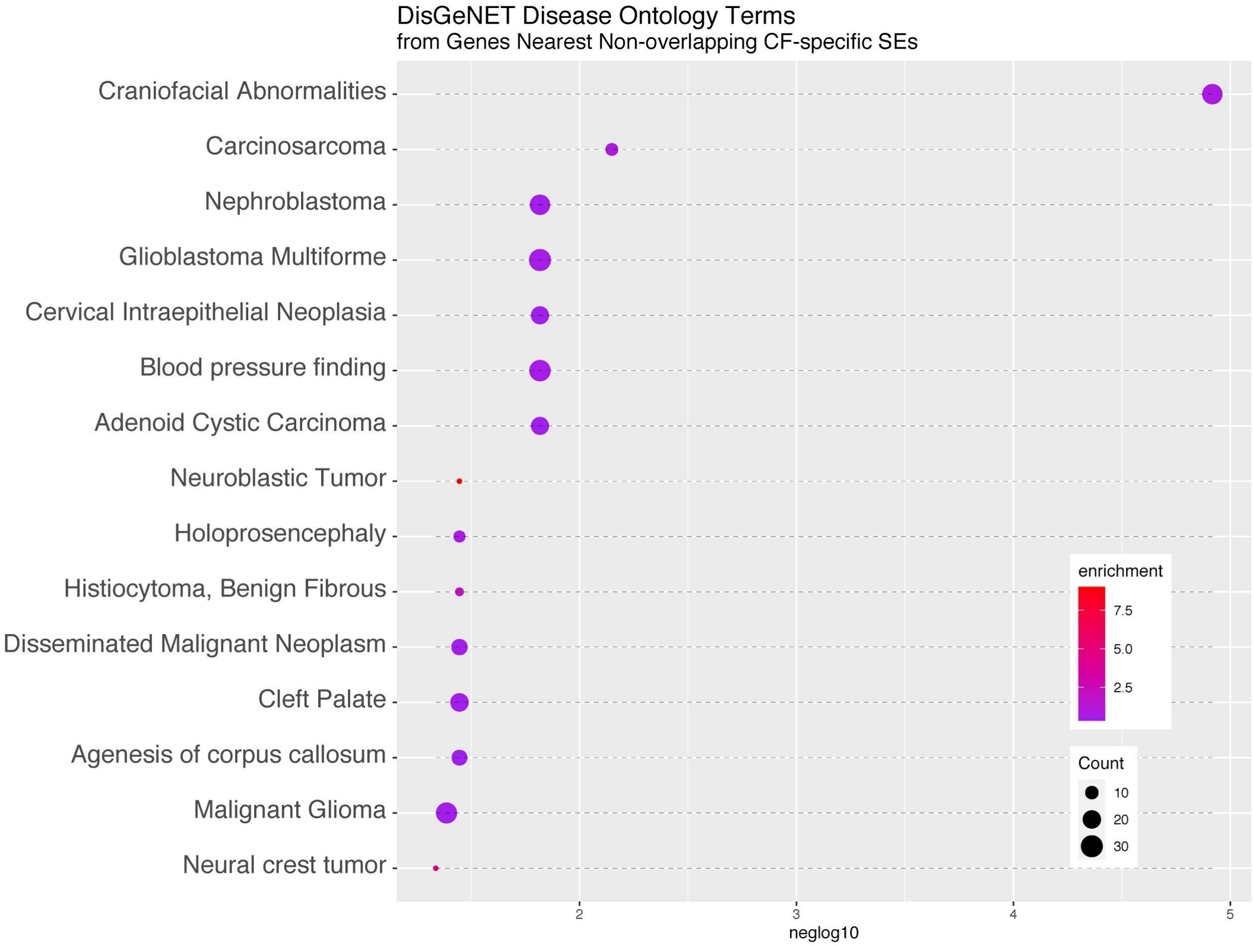


**Figure 1 Supplement 3-** Disease Ontology terms from DisGeNet enriched in genes assigned to superenhancers in non-coding regions. Assignment of the two nearest genes was done through the BEDTools suite function "closest".

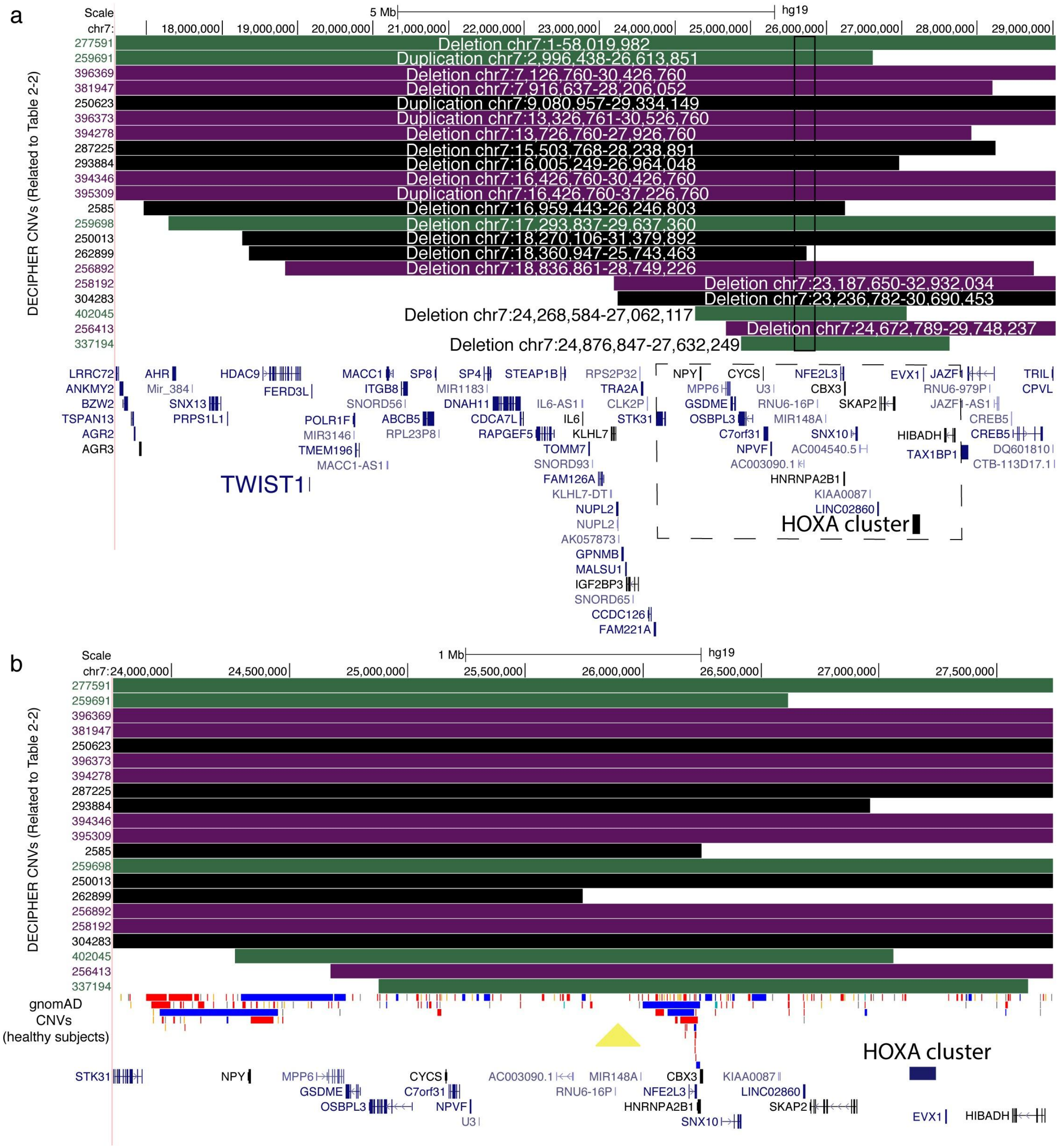


**Figure 2 Supplement 1-** a. Copy number variants from DECIPHER database overlapping the putative novel craniofacial superenhancer region (black box with solid outline). CNVs represented by green bars have a noted phenotype but do not include a specifically described craniofacial phenotype, purple bars have a specifically described craniofacial phenotype and black bars have no phenotype reported. b. Enlargment of region in box with dotted outline.The track for gnomAD CNVs, filtered for CNVs >300bp appears below the DECIPHER CNV bars, blue represents gains and red losses. 300bp based on typical size range of CNVs identified in healthy human populations (Zarrei et al., 2015). A region with a notable lack of CNVs in gnomAD subjects is marked by a yellow triangle.

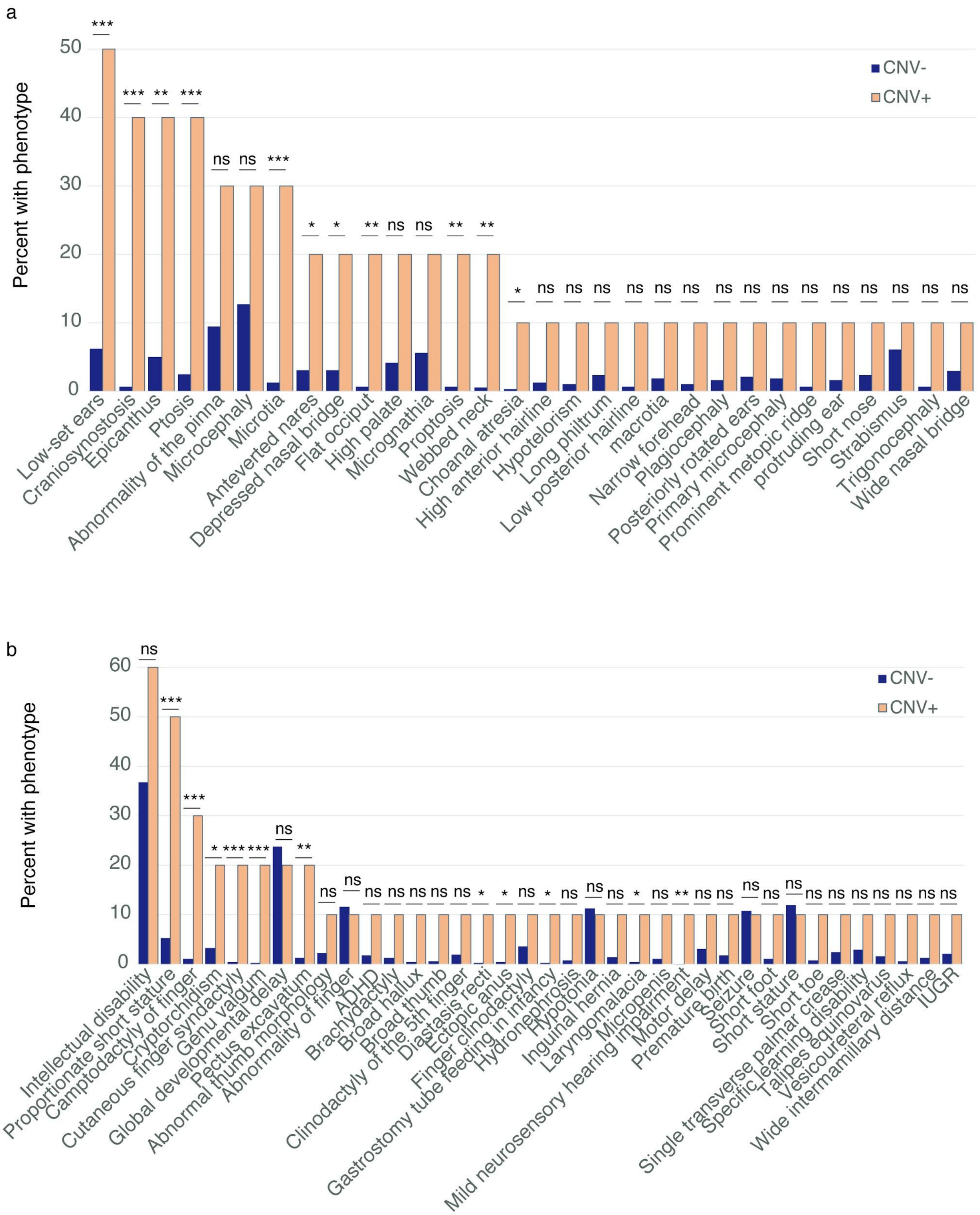


**Figure 2 Supplement 2-** Frequency of phenotypes present in individuals within the DECIPHER Database with CNVs overlapping chr7: 25,580,400-25,849,400 compared to frequency of those phenotypes in the

the DECIPHER Database for CNVs not overlapping the region. Statistical test is Fisher Exact Test. * p<0.05, ** p<0.01, *** p<0.001

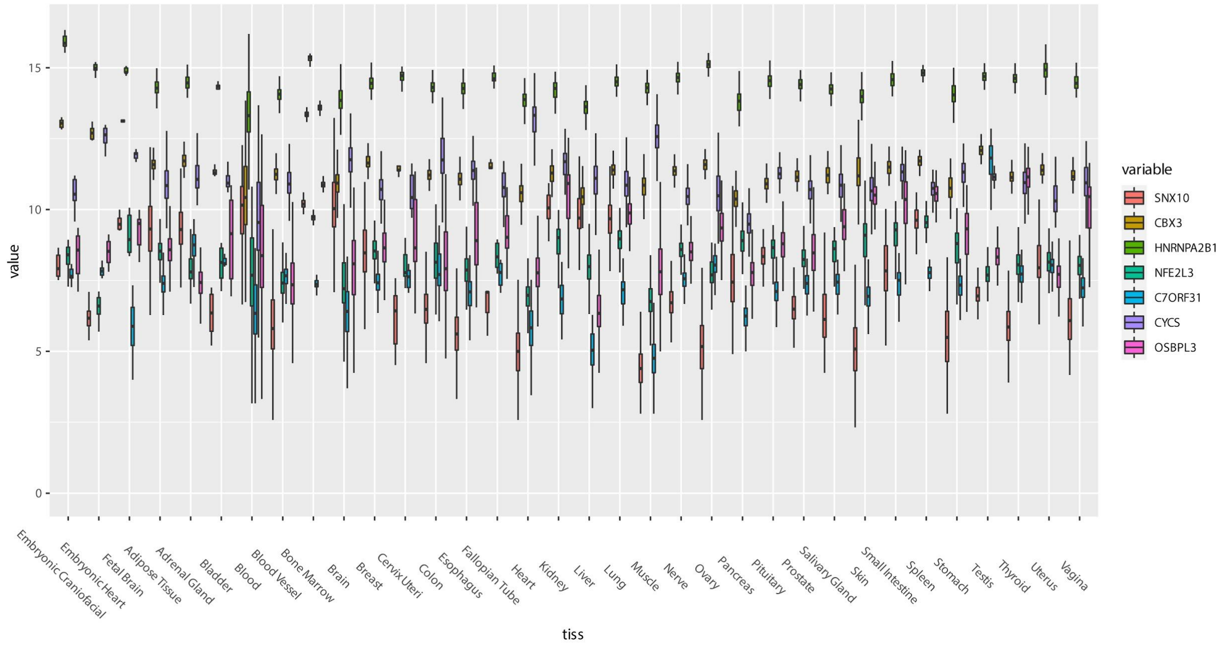


**Figure 2 Supplement 3- Expression of genes within 500kb of gene desert.**

The $\log_2 + 1$ of counts from human primary craniofacial tissue, embryonic heart, fetal brain and 31 adult tissues from GTEx. These genes show ubiquitous expression across all tissues surveyed.

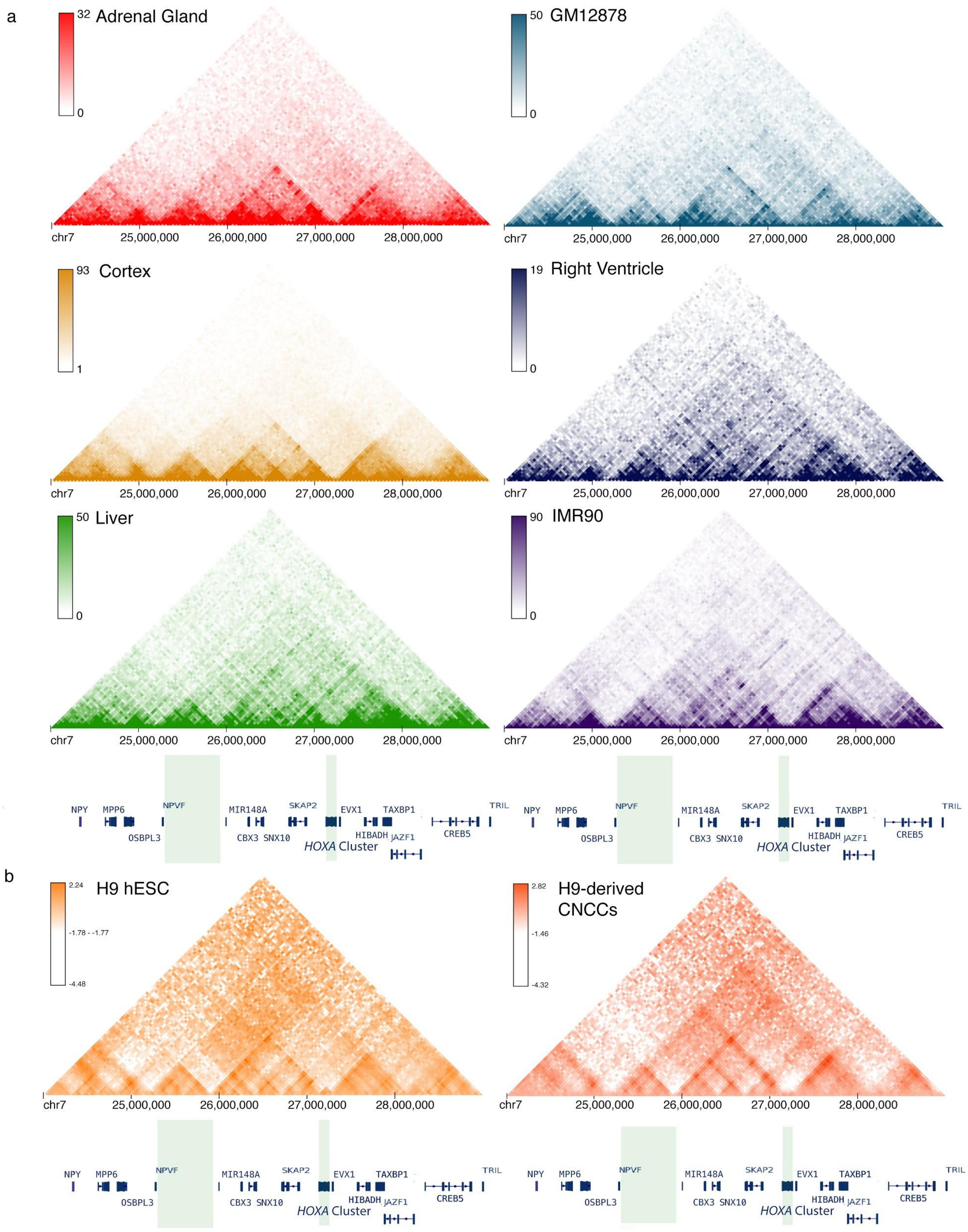


**Figure 3 Supplement 1-** HiC from publicly available data of various human tissues and cell lines (a) and publicly available data from H9 (b- left) and H9-derived CNCCs (this study, b-right). Plots were

generated through the HiC Browser hosted by Northwestern University. *HOXA* cluster and putative novel superenhancer region are highlighted with green shading.

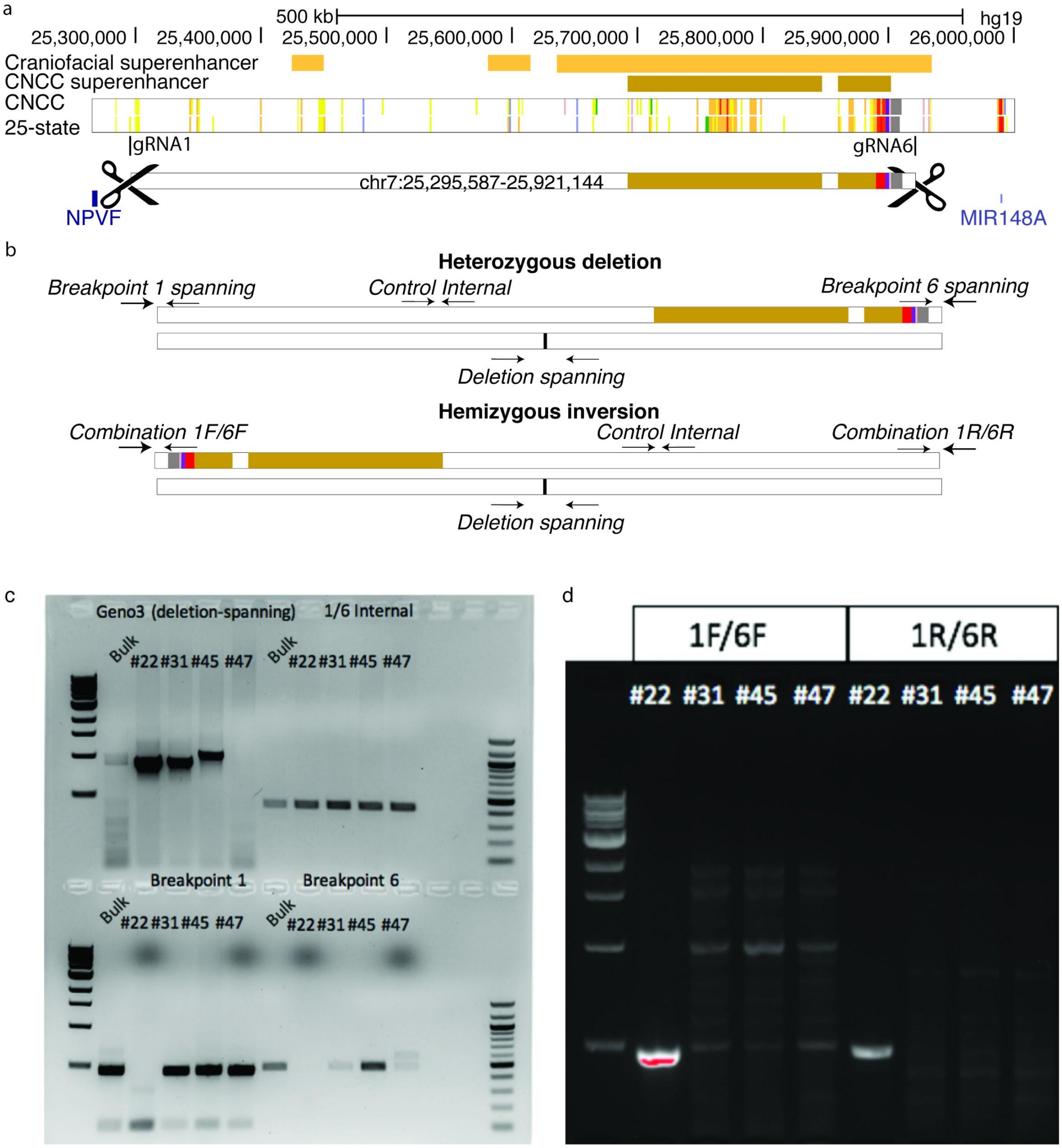


**Figure 4 Supplement 1-** a. Location of guide RNAs gRNA1 and gRNA6 relative to the WT orientation. b. Screening strategy for determining whether clones are heterozygous for the 1/6 deletion and determining if a clone contains an inversion of the targeted region. c-d. PCR results identifying heterozygous clones and the clone carrying the hemizgous inversion (#22, also referred to as INV).

**Figure 4 Supplement 2-** (Word document) Sequence across breakpoints for clones #31 (heterozygous) and #22 (inversion, INV). Clone sequence based on Sanger sequencing results (see Methods) and compared against hg19 assembly.

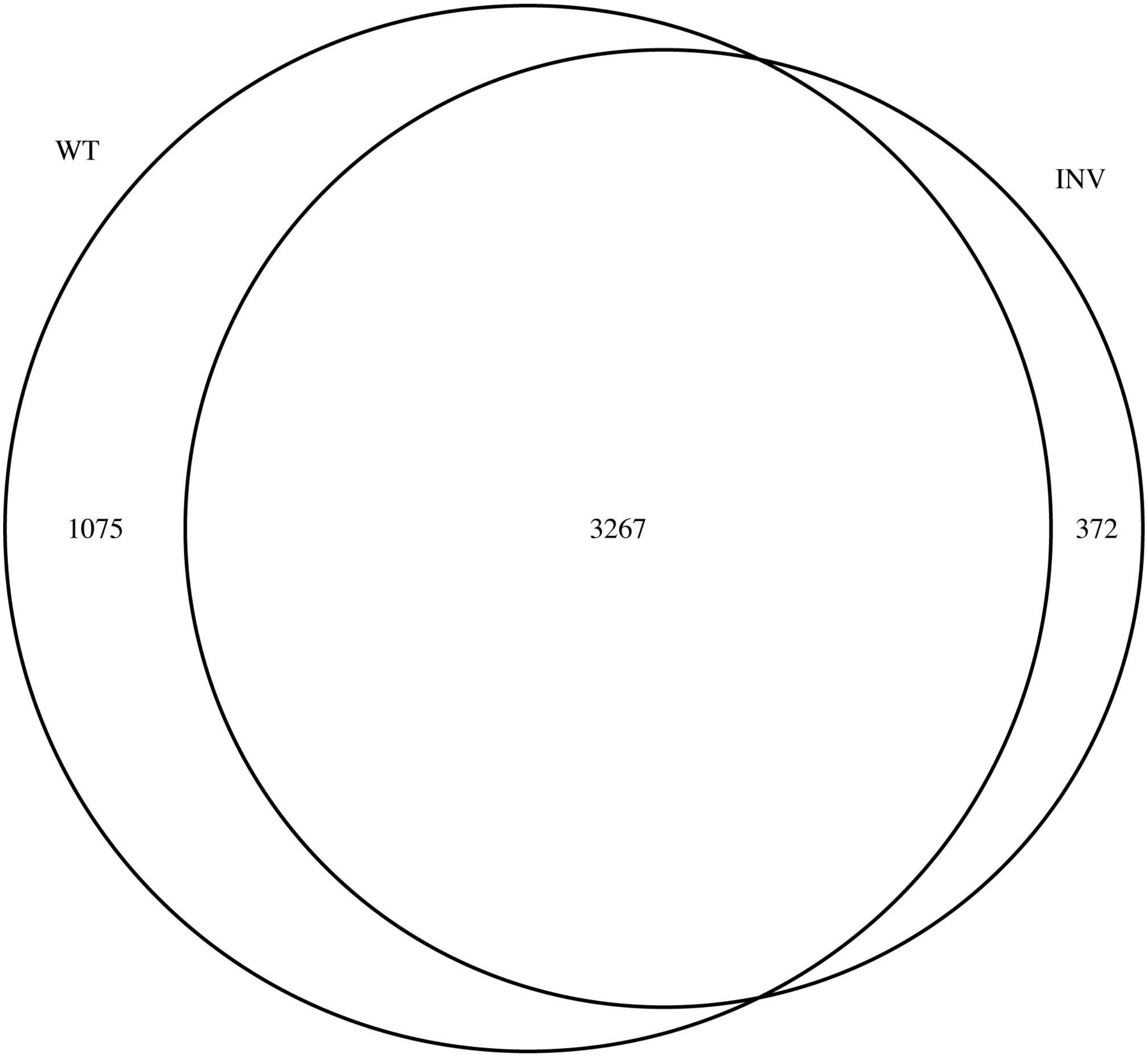


**Figure 4 Supplement 3-** Venn diagram of genes expressed in WT and INV H9 lines at d0 of CNCC differentiation (baseline).

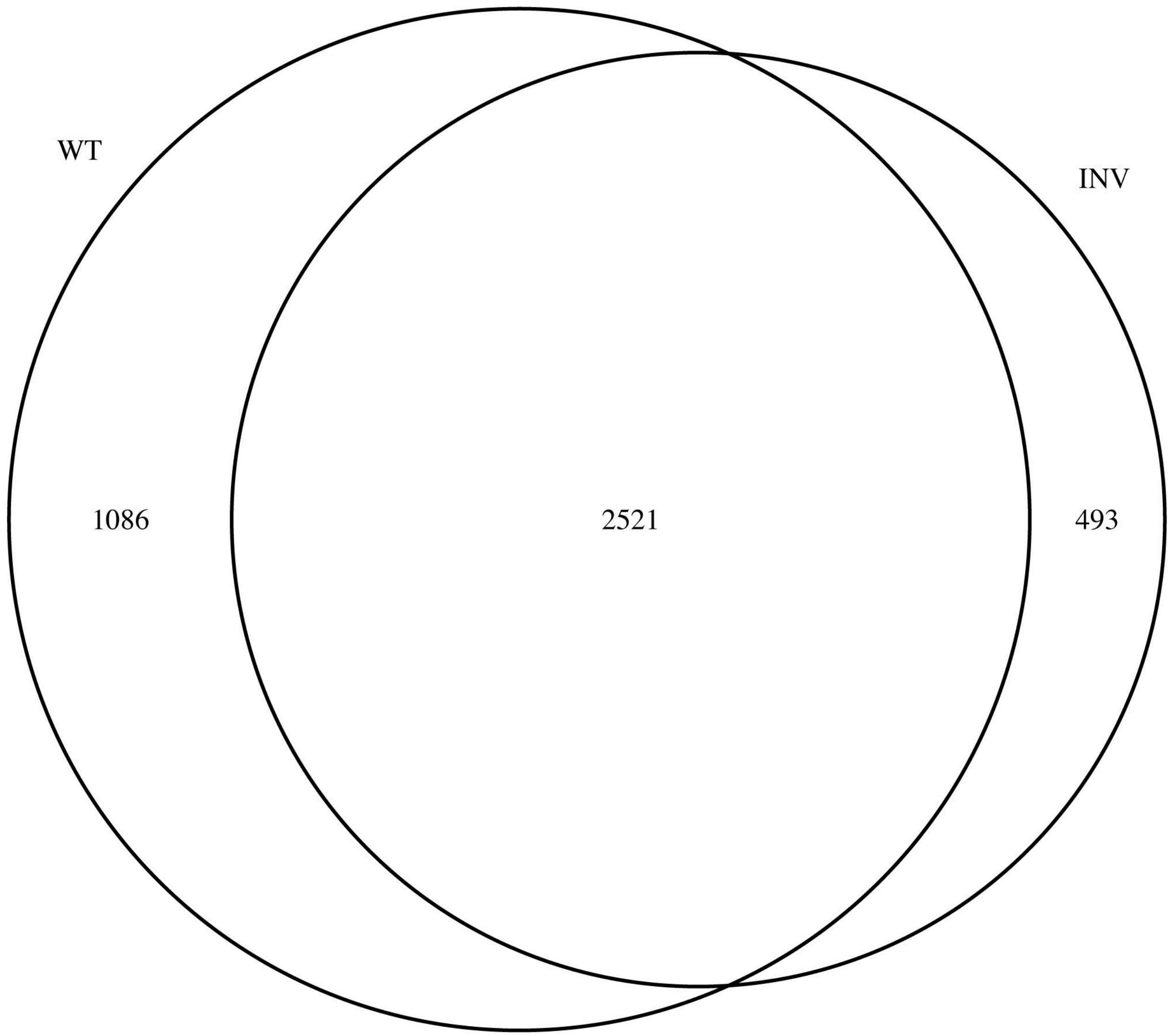


**Figure 4 Supplement 4-** Venn diagram of genes expressed in WT and INV H9 lines at d5 of CNCC differentiation.

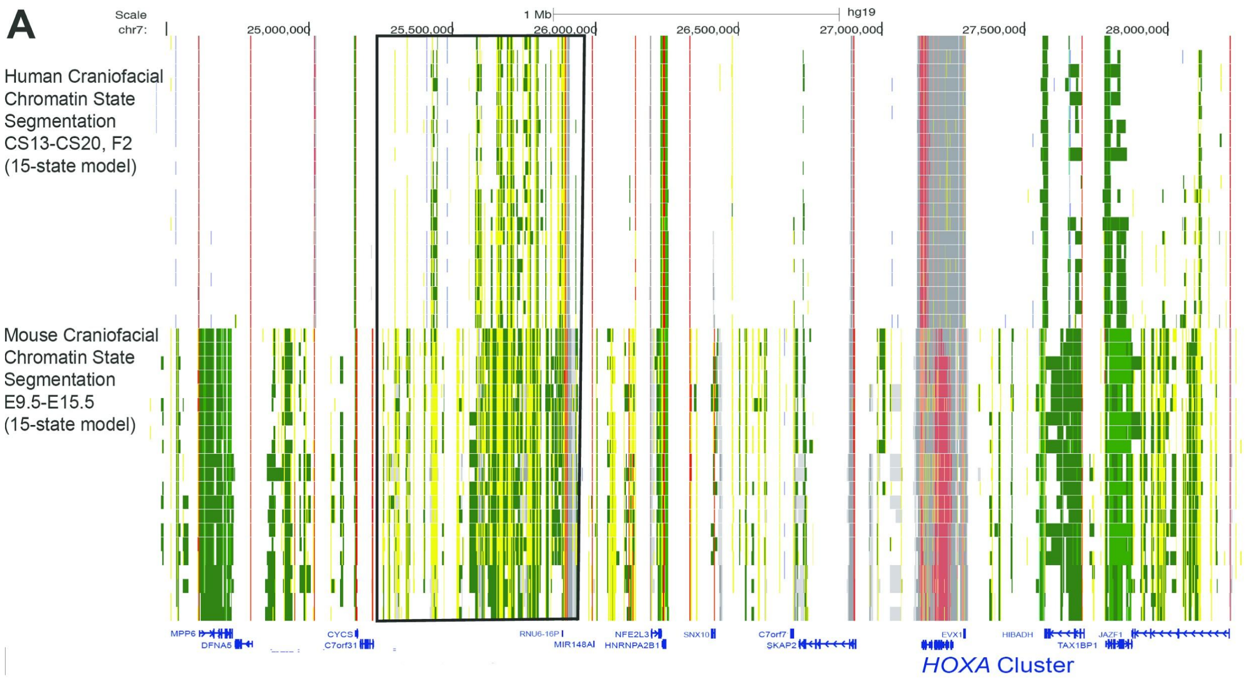


**Figure 5 Supplement 1-** Comparison of chromatin states in the 15-state model between human embryonic craniofacial tissue (hg19) and mouse embryonic craniofacial tissue (mm9 lifted to hg19).

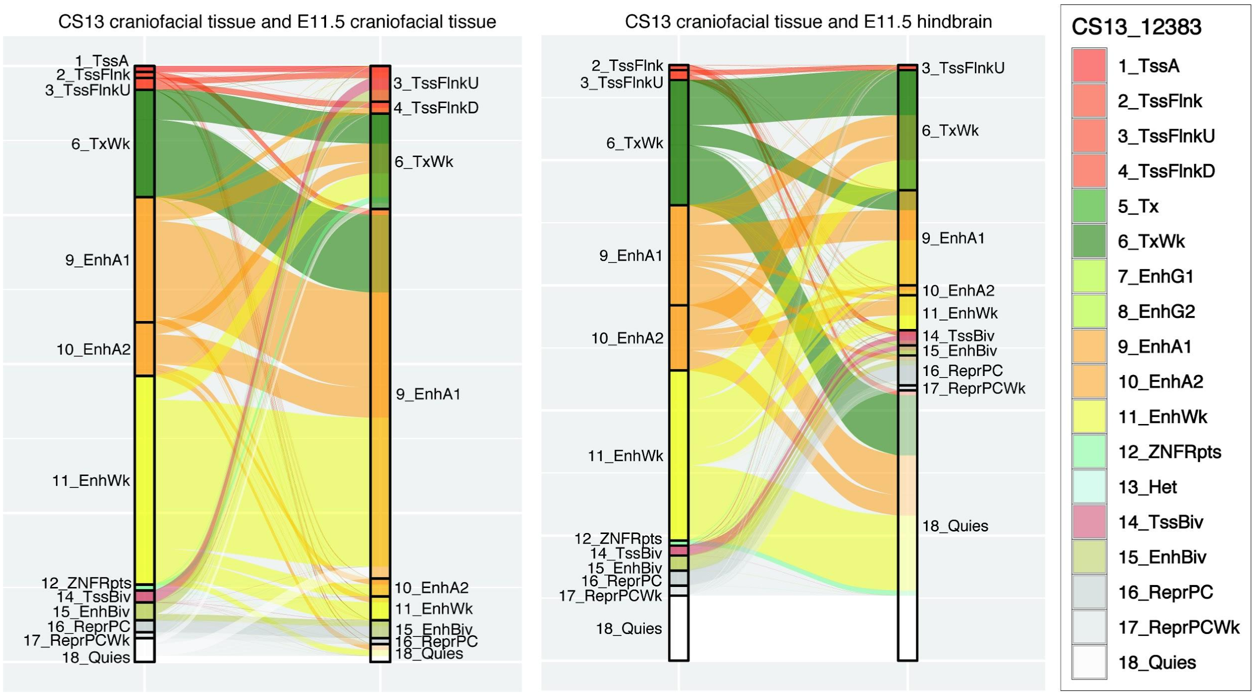


**Figure 5 Supplement 2-** Chromatin state composition comparison for 18-state model between CS13 craniofacial tissue and E11.5 craniofacial tissue or E11.5 hindbrain. Mouse chromatin states lifted from mm10 to hg19 using liftOver.

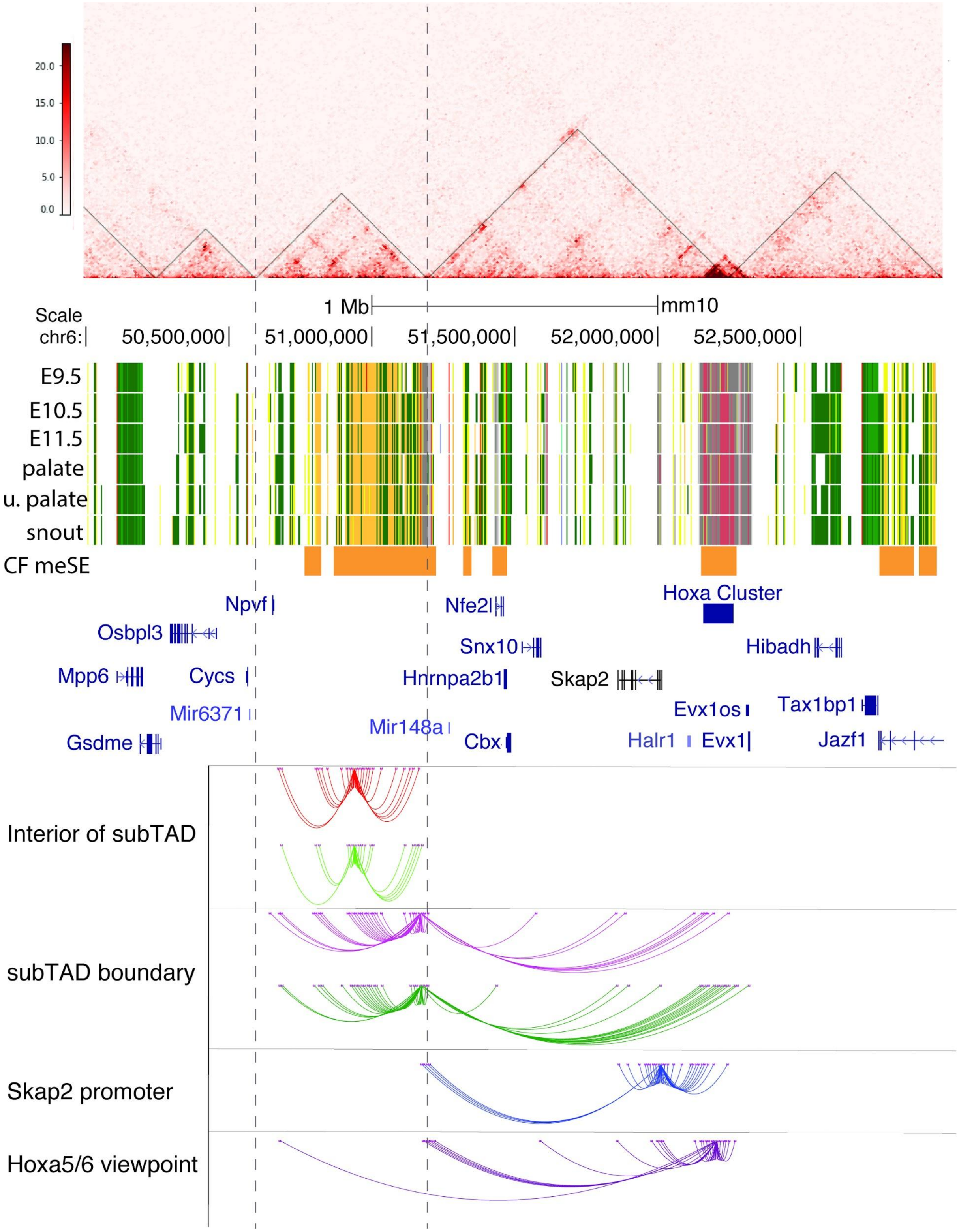


**Figure 5 Supplement 3-** HiC data from WT E11.5 mouse craniofacial tissue, TADs or subTADs determined at 50Kb resolution. The superenhancer subTAD is marked throughout with dotted lines (Top). Chromatin states in 18-state model and mouse embryonic craniofacial superenhancers shown in orange bars above gene notations (Middle). Interactions identified though 4C-seq at all viewpoints tested

(Bottom). Viewpoint1(red) and Viewpoint 2 (light green) are located within the interior of the superenhancer subTAD. Viewpoint 3 (magenta) and Viewpoint 4 (dark green) are located at the boundary of the subTAD. Viewpoint 5 (dark blue) is located near the *Skap2* promoter and Viewpoint 6 (purple) is located near the intergenic space between *Hoxa5* and *Hoxa6*.

**Figure 6 Supplement 1-** (Separate Large File) Ventral views of the cranial base from multiple WT, ΔGCR/+ (HT) and ΔGCR/ΔGCR (KO) E18.5 mice.

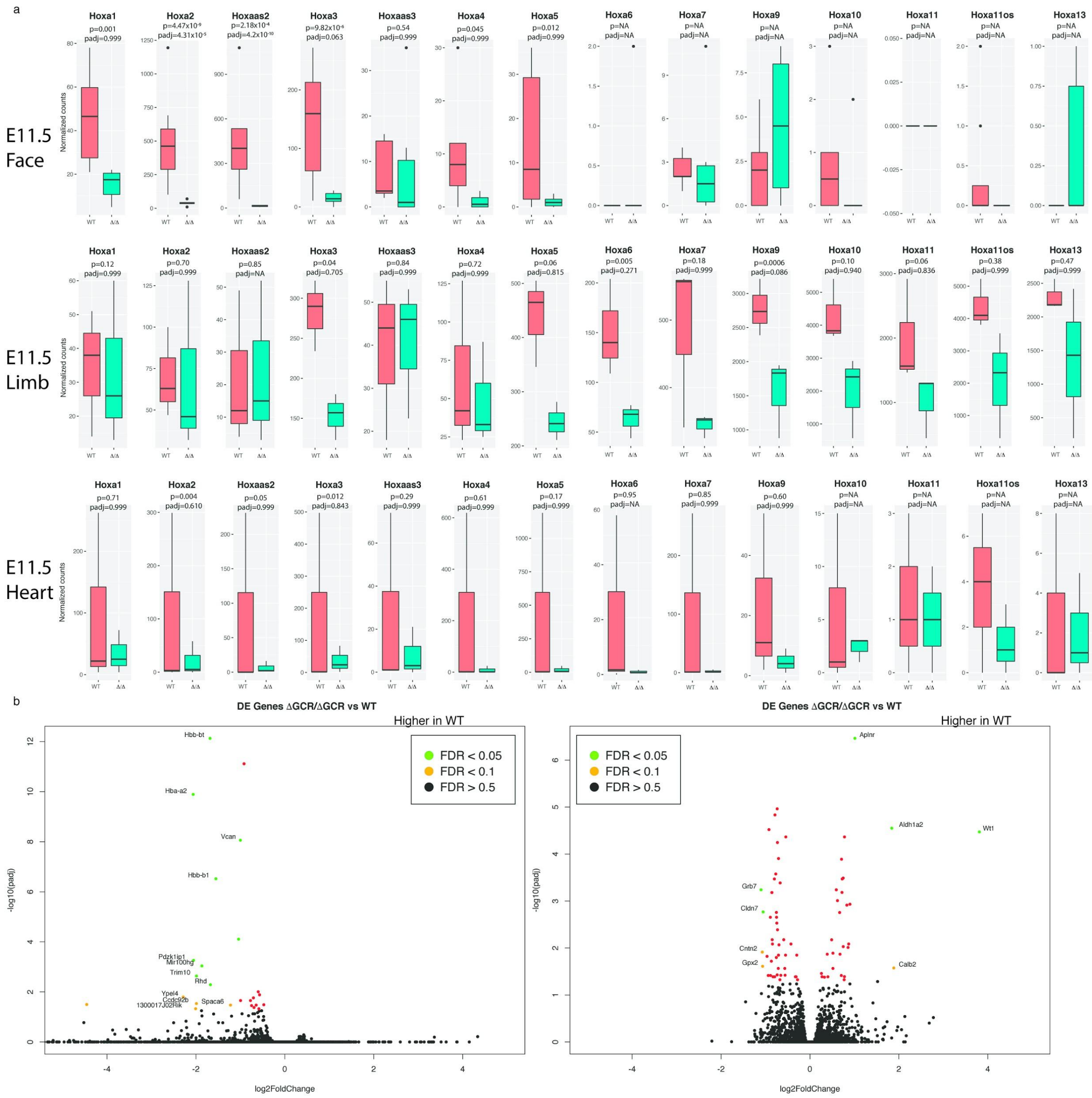


**Figure 7 Supplement 1-** Comparison of *Hoxa* gene expression in embryonic face, limb and heart (a). Gene expression changes in heart (b-left panel) and limb (b-right panel). Few significant genes change as a result of the deletion, and did not include the *Hoxa* genes in either heart or limb.

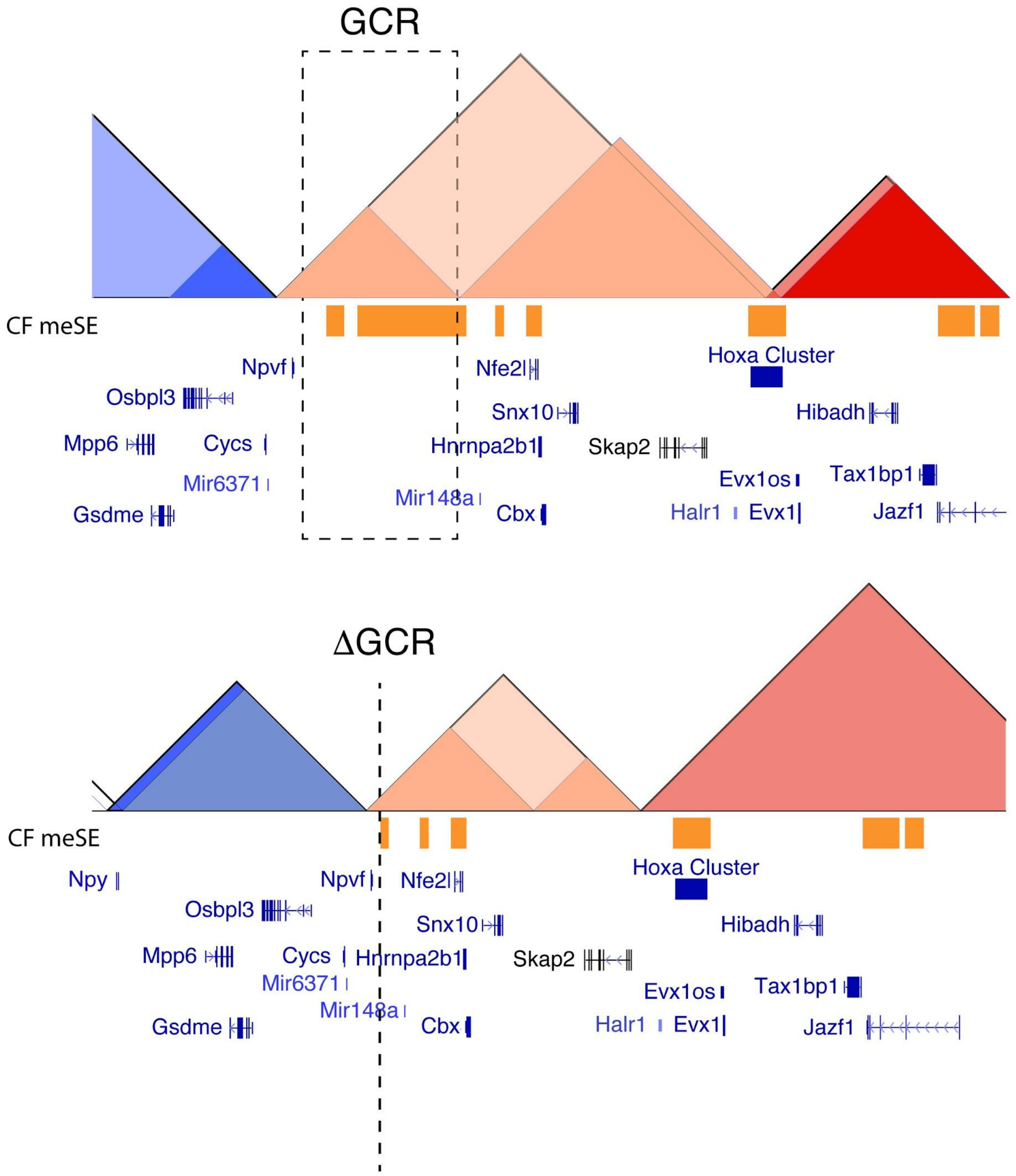


**Figure 7 Supplement 2-** a. Deletion of GCR re-structures the *HoxA* regulatory landscape. The removal of ~625kb and almost complete removal of the superenhancers in the gene desert between *Npvf* and *Mir148a* disrupts the contact between the GCR and the *HoxA* cluster. Schematics based on HiC of E11.5 cranofacial tissue from WT (top panel) and △GCR/△GCR mice (bottom panel). The bottom panel was created by alignment to a custom genome based on mm10 with deletion of the GCR coordinates. Presentation of superenhancers in the bottom panel are predicted, based on the superenhancers as they appear in the WT. TADs predicted at 100Kb are represented with black outline, TADs predicted at 50Kb have no outline.

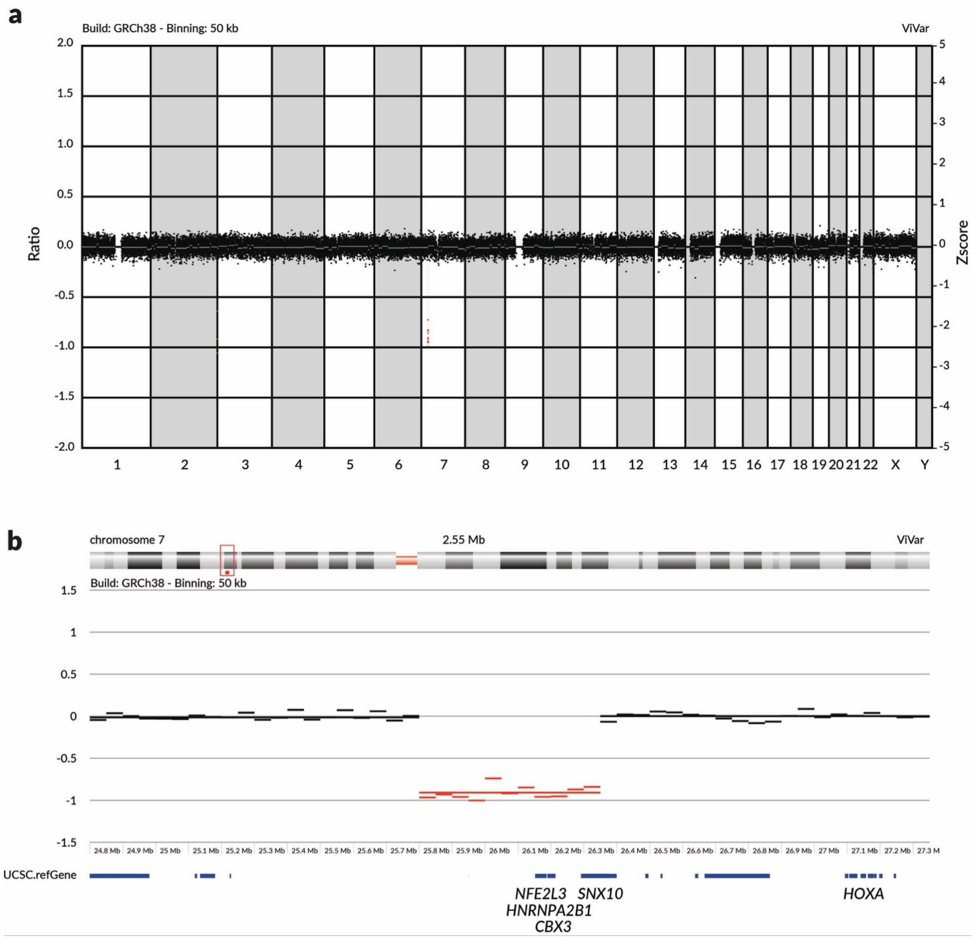


**Figure 8 Supplement 1- Identification of a 550kb *de novo* deletion upstream of the *HOXA* cluster through shallow whole genome sequencing and copy-number variant (CNV) analysis using ViVar.** (a) whole genome lineview. (b) zoomed in on deletion (binning: 50kb). ViVar analysis and visualization (Sante et al., 2014).

### Supplemental References

Sante, T., Vergult, S., Volders, P.-J., Kloosterman, W.P., Trooskens, G., De Preter, K., Dheedene, A., Speleman, F., De Meyer, T., and Menten, B. (2014). ViVar: a comprehensive platform for the analysis and visualization of structural genomic variation. PLoS One *9*, e113800.
